# Structure of a Rad52 homolog from bacteriophage in complex with a novel duplex intermediate of DNA annealing

**DOI:** 10.1101/2022.03.17.484533

**Authors:** Brian J. Caldwell, Andrew Norris, Vicki H. Wysocki, Charles E. Bell

## Abstract

Human Rad52 protein binds to ssDNA and promotes the annealing of complementary strands. This activity is central to multiple DNA repair pathways and Rad52 is a target for cancer therapeutics. Previous crystal structures of the DNA binding domain of Rad52 revealed an 11-mer ring that binds to ssDNA in an extended conformation with the bases exposed for homology recognition. While this complex is likely involved in the early stages of annealing, there is no structure of Rad52 with two strands of DNA bound simultaneously, and its mechanism of annealing is poorly understood. To approach this problem, we have turned to the RecT/Redβ family of annealing proteins from bacteriophage, which are distant homologs of Rad52 that form stable complexes with a duplex intermediate of annealing. We have used single particle cryo-electron microscopy (cryo-EM) to determine a 3.4 Å structure of a RecT homolog from a prophage of *Listeria innocua* (LiRecT) in complex with two complementary 83-mer oligonucleotides that were added to the protein sequentially. The structure reveals a left-handed helical filament of the protein bound to a novel conformation of DNA duplex that is highly extended and under-wound. The duplex is bound at a stoichiometry of 5 bp/monomer to a deep, narrow, positively-charged groove that runs along the outer surface of the filament. Data from native mass spectrometry confirm that the filament complex seen by cryo-EM also exists in solution. Collectively, these data provide new insights into the mechanism of annealing by LiRecT and by homologous proteins including human Rad52.

## Introduction

Human Rad52 (hRad52) protein binds to single-stranded DNA (ssDNA) and promotes the annealing of complementary DNA strands (1). This reaction is involved in multiple aspects of genome maintenance, including two of the three main pathways for repair of double-stranded DNA (dsDNA) breaks (2,3), recovery of stalled replication forks (4,5), and extension of telomeres (6). Although not required for viability of normal human cells, mutations in Rad52 are synthetically lethal with loss of DNA repair factors that are frequently mutated in cancer, most notably BRCA1 and BRCA2 (7,8). This makes pharmacological inhibition of Rad52 a viable strategy for treatment of these cancers in the same way that poly(ADP-ribose)-polymerase inhibitors have been exploited. Since this discovery, several groups have developed small molecule inhibitors of Rad52, primarily by high-throughput screening of compound libraries against fluorescence-based DNA binding and annealing assays (9). By contrast, rational design of Rad52 inhibitors based on a fundamental understanding of its mechanism has yet to be fully exploited.

Crystal structures of the N-terminal DNA binding domain of Rad52 revealed an 11-mer ring with two distinct DNA binding sites (10–12). One site contains a 40-nt ssDNA bound at the base of a deep, narrow, positively-charged groove that wraps around the upper surface of the ring. This DNA is bound in an extended conformation with its bases exposed for homology recognition (12). The second site, identified in a separate crystal structure, lies at the upper rim of the same DNA binding groove. This site contains a shorter segment of ssDNA bound in a right-handed helical conformation that bridges two neighboring rings of Rad52 in the crystal lattice (12). The role of this second site is not yet fully understood, but mutational analysis indicates that it is required for annealing (13). While it seems clear that the first structure represents a substrate complex of the protein with the first DNA, there is no structure of Rad52 bound with two complementary strands of DNA simultaneously, and its overall mechanism of annealing is poorly understood.

To address this problem, we have turned to a group of annealing proteins from bacteriophage that share limited homology with Rad52 (14). These proteins, which are annotated as the RecT/Redβ superfamily based on the members found in *E. coli* and bacteriophage λ, offer a potential advantage in that they bind with high affinity to a stable intermediate of annealing formed when two complementary DNA strands are added to the protein sequentially (15). Negative stain EM of Redβ from bacteriophage λ (λ-Redβ) revealed that it forms oligomeric rings like Rad52 when bound to ssDNA, but left-handed helical filaments when bound to annealed duplex (16). These data led to a model for annealing in which the ring form of Redβ binds to ssDNA with its bases exposed for homology recognition, while a helical filament of the protein assembles on the duplex intermediate as it is formed to drive the annealing reaction forward. However, due to the low resolution of these structures, the locations of the DNA-binding sites and the details of the protein-DNA interactions could not be resolved.

Here, we have used single-particle cryo-EM to determine a 3.4Å structure of a homolog of λ-Redβ from the A118 prophage of *Listeria innocua* that we will call LiRecT. The structure reveals a left-handed helical filament of the protein bound to a novel 83-mer duplex intermediate of annealing. The filaments are strikingly similar to those seen for λ-Redβ at low-resolution over twenty years ago, but our structure now reveals the fold of the protein, the location of the DNA binding groove, the conformation of the DNA, and the details of the protein-DNA and inter-subunit interactions.

## Results

### Architecture of the LiRecT-DNA Complex

The LiRecT protein was purified and found to bind to ssDNA and form a complex with annealed duplex in a similar manner as λ-Redβ (Fig. S1). For cryo-EM analysis, a complex of LiRecT with annealed duplex was formed by incubating the protein with two complementary 83-mer oligonucleotides that were added to the protein sequentially. The sequences of the oligonucleotides were derived from a naturally occurring sequence in M13 DNA described previously (15,17). The complex appeared as helical filaments of varying lengths, including some with endon views (Fig. S2). Standard single-particle analysis without helical symmetry averaging in cryoSPARC yielded a 3.4Å reconstruction with fully interpretable density for the LiRecT subunits and the bound DNA at the central portion of the filament (Fig. 1).

**Fig. 1.**
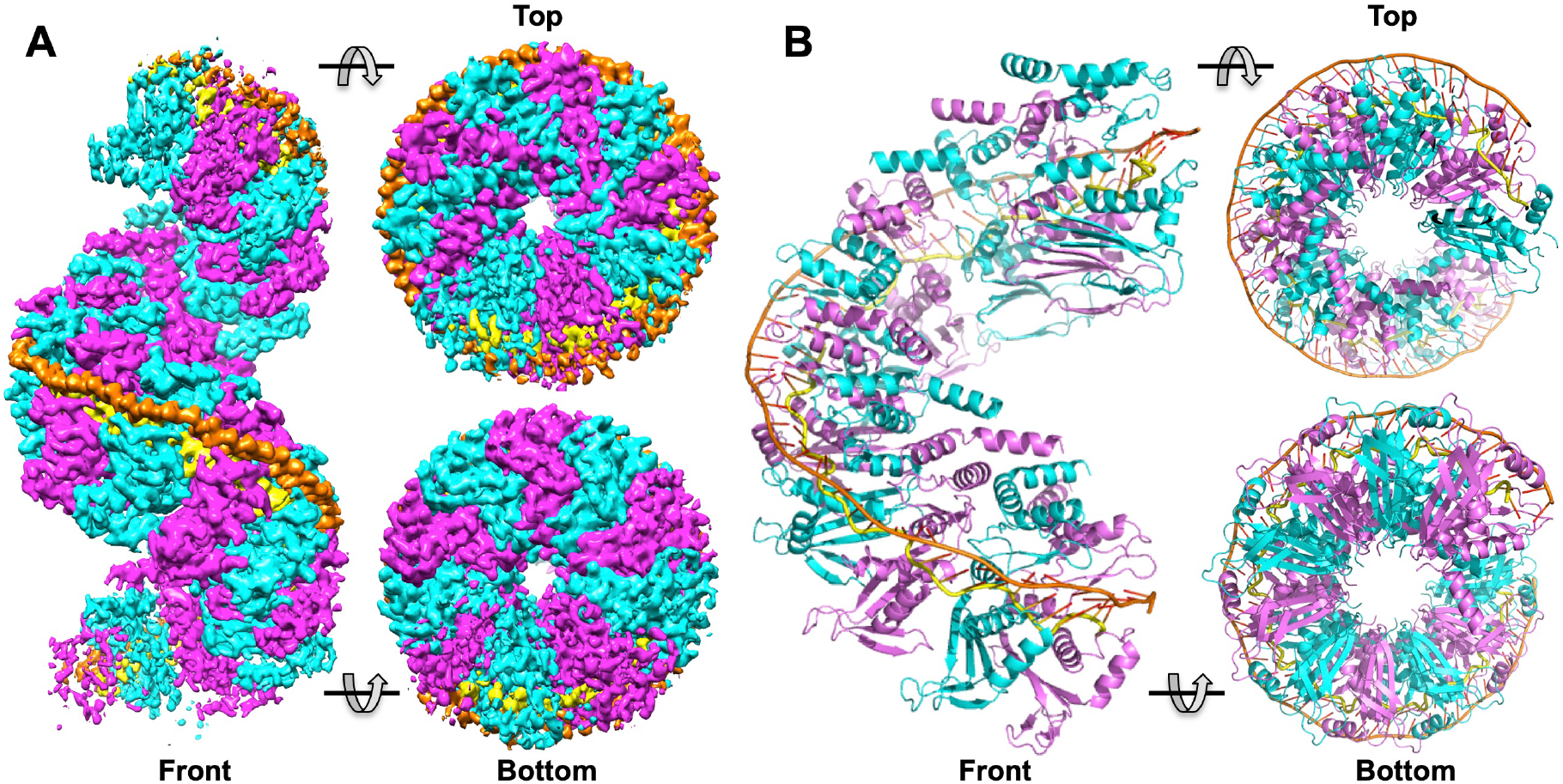
Cryo-EM structure of LiRecT bound to 83-mer annealed duplex. (***A***) Views of the full 18-subunit 3D reconstruction. Alternating subunits of LiRecT are colored cyan and magenta. The inner DNA strand is yellow and the outer strand is orange. Notice that the density gets progressively weaker towards the ends of the filament. (***B***) Ribbon diagrams of a 10-subunit model fit and refined to the density for the central portion of the filament. Notice that the duplex is bound in a novel conformation that is highly extended and under-wound.

In the complex, LiRecT assembles into a left-handed helical filament that is highly reminiscent of those seen previously for λ-Redβ (16). The filament has an open corkscrew-like shape with an inner diameter of 20 Å, an outer diameter of 100 Å, and a pitch of 105 Å with approximately 10 subunits per turn. The two complementary 83-mer strands are bound as a highly extended and under-wound duplex to a deep, narrow, positively-charged groove that runs along the outer surface of the filament (Figs. 1 and S3). One strand, which we call “inner” (yellow in Fig. 1), is bound to the deepest part of the groove with its nucleotide bases facing outwards. The other strand, which we call “outer” (orange), is bound to the outer portion of the groove with its bases facing inward to form normal Watson-Crick base pairs with the inner strand.

Each monomer of LiRecT binds to 5 bp of DNA. Based on this ratio, we would expect the filament to contain 16-17 subunits of LiRecT bound to the 83-mer duplex. While we do see a filament of approximately this length in the 3D reconstruction, the density towards the ends of the filament gets progressively weaker, presumably due to flexibility and/or imperfect alignment of the particles along the filament axis. Consequently, we chose to refine a model that consists of just the 10 subunits of protein and 48 bp of DNA at the central portion of the filament (Fig. 1*B*), for which the density is strongest. Due to the helical symmetry however, this model likely encompasses all of the relevant protein-protein and protein-DNA interactions that exist in the full filament, except at the ends. In addition, although the resolution of the map was high enough to clearly see nucleotide bases (Fig. S2*I*), purines and pyrimidines could not be distinguished, likely due to the imperfect alignment of the particles along the filament axis. The DNA has thus been modeled as dT48 for the outer strand, and dA48 for the inner strand, despite the fact that both strands contain a natural variation of all four nucleotides. Finally, based on the measured helical parameters of 10 subunits per turn, we would expect the filaments to contain approximately 1.5 turns. In the cryo-EM images however, many of the filaments contain several turns (Fig. S2*A*), suggesting that they can stack end-to-end. The result of single particle analysis however converged on just a single 1.5-turn filament.

### LiRecT Monomer Fold and Relation to Rad52

The structure reveals that LiRecT and by extension the RecT/Redβ family of proteins does indeed share homology with Rad52, as had been predicted (18–20). The two structures superimpose to an RMSD of 4.3 Å for 107 pairs of Cα atoms that share 14% sequence identity. This common core covers 56% of the LiRecT structure of 191 amino acids, and 40% of the full-length LiRecT protein of 271 amino acids. Despite this common core, Rad52 was not identified as a hit in a DALI search for structural homology on LiRecT (21), reflecting the high degree of structural divergence. The common core fold consists of 2 central α-helices (α2 and α3) that form the base of the DNA binding groove, combined with a beta hairpin (β1-β2) on one side and a three-stranded antiparallel beta sheet (β3-β5) on the other. In Fig. 2, we have numbered the common core secondary structural elements of LiRecT based on Rad52, and used letters for inserted elements, which are shown in green. The first insertion is an N-terminal 3-helix bundle (αA, αB, αC) that sits at the upper rim of the filament and packs with neighboring copies of itself from the adjacent subunits (Fig. 3). The second is a β-hairpin (βA-βB) inserted after β3 that interacts with β3 of the neighboring subunit at the lower rim of the filament (Fig. 3). The third is a pair of α-helices (αD and αE) inserted after β5 that pack with α3 to form the lower rim of the DNA-binding groove (Fig. 3). The modeled portion of each LiRecT monomer consists of residues 34-224 of the 271 amino acid protein. The additional residues at the N- and C-terminal ends, which are presumably disordered relative to the main body of the filament, would project from the upper and inner surfaces of the filament, respectively (Fig. 3A). Comparisons of the LiRecT structure to predicted structures of it and of λ-Redβ from RoseTTAFold are shown in Fig. S4 (22). A structure-based sequence alignment of LiRecT and λ-Redβ is shown in Fig. S5.

**Fig. 2.**
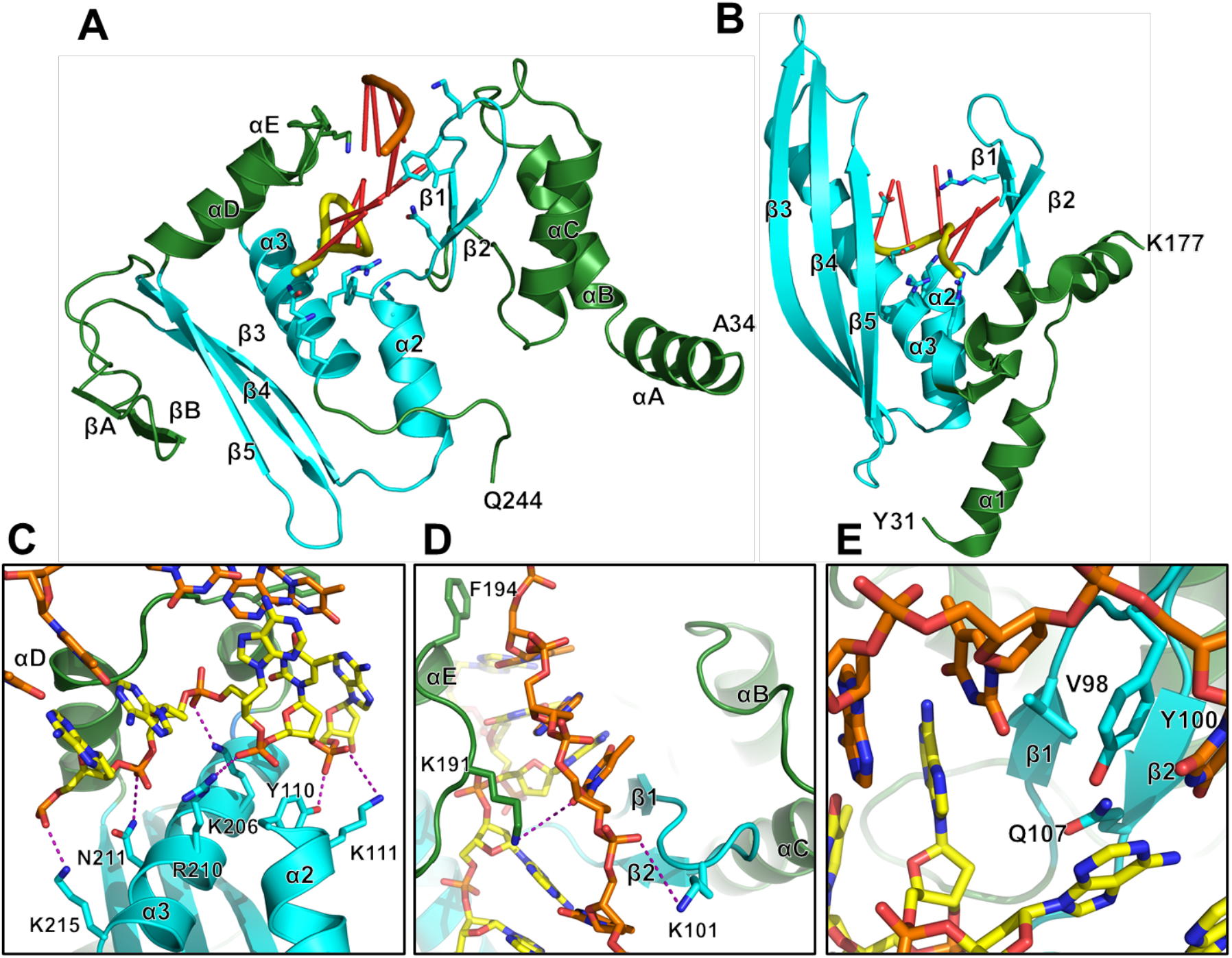
Structure of the LiRecT monomer, comparison with Rad52, and interactions with DNA. (***A,B***) Monomers of LiRecT (A) and Rad52 (B) are shown with their common core fold in cyan and extraneous segments in green. The DNA binding groove is formed by the 2 central α-helices (α2, α3), the β1-β2 hairpin, and αE-αD (LiRecT) or β3-β5 (Rad52). Rad52 is drawn with coordinates from PDB accession ID 5XRZ (12). (***C,D,E***) Close-up views of LiRecT interactions with the inner strand (C), outer strand (D), and β1-β2 hairpin (E). Hydrogen bonds within 3.5Å are shown as dotted lines.

**Fig. 3.**
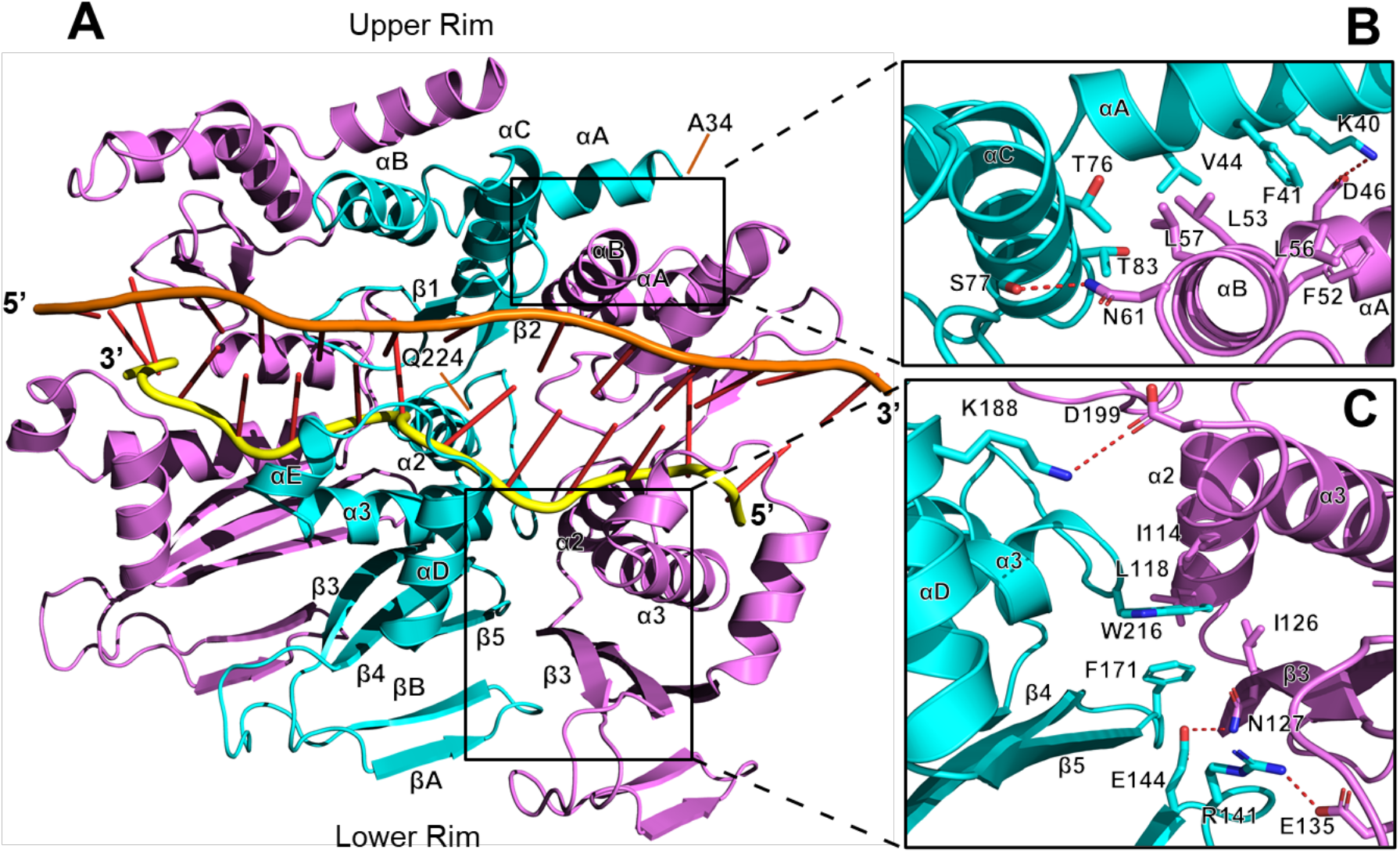
Close-up view of the DNA binding groove and inter-subunit packing. (***A***) Front view of 3 subunits of the LiRecT filament with secondary structures and terminal residues (A23 and Q224) labeled for the middle (cyan) subunit. (***B,C***) Close up views of the inter-subunit interactions above (B) and below (C) the DNA binding groove. Hydrogen bonds within 3.5Å are shown as dotted lines.

### Protein-DNA Interactions

The two strands of DNA are bound to a deep, narrow, positively-charged groove that runs along the outer surface of the filament. The base of the groove is formed by α2 and α3 and its outer walls are formed by the β1–β2 hairpin on one side and the αD–αE insertion on the other (Fig. 2 & 3). The inner strand is bound to the deepest part of the groove where it is contacted by the side chains of Y110 and K111 from α2, and K206, R210, N211, and K215 from α3 (Fig. 2C). These residues form extensive interactions with the sugar phosphate backbone of the inner strand and hold it in an irregular conformation that is periodically kinked (Fig. 3). By contrast, the outer strand makes far fewer interactions with the protein and adopts a smoother conformation that is held in place primarily by normal Watson-Crick base pair interactions with the inner strand. The few residues that do contact the outer strand are K101 at the tip of the β1-β2 hairpin, and K191 and F194 from the αD–αE insertion (Fig. 2D). Although most of the contacts involve the sugar-phosphate backbone of each strand, the side chains of V98, Y100, and Q107 from the β1–β2 hairpin wedge into the base pairs at every 5^th^ bp step to separate them (Fig. 2E). Specifically, the phenyl ring of Y100 of each subunit stacks with the base of every 5^th^ nucleotide of the outer strand, while the side chain of Q107 stacks with the opposing base of the inner strand. These interactions introduce a dramatic kink in the backbone of the inner strand where the bases are separated (Fig. 3). Most of the residues that contact the DNA, particularly those that contact the inner strand, are highly conserved among six distant homologs of LiRecT identified by PSI-BLAST (Fig. S6). This suggests that the structure has captured a functionally relevant state of the protein.

Although the two DNA strands mostly contact one another via normal Watson Crick base pair interactions, the duplex is highly extended and under-wound compared to B-form DNA (Fig. S7 and S8). In concert with the 5 bp/monomer stoichiometry, the bases are stacked in a repeating pattern, with groups of 5 bp stacked with approximately 3.8 Å spacing, alternating with a larger 9 Å spacing where the β1–β2 insertion occurs. Overall, the duplex is about 1.5 times as extended as B-form DNA and is completely unwound. The local base pair step parameters deviate significantly from B-form DNA in a regularly repeating manner every 5 bp (Fig. S8). This is largely due to the irregular and bent conformation of the inner strand.

### Inter-subunit Packing

The LiRecT subunits pack in the filament with interactions that bury 1830 Å^2^ of total solvent accessible surface area. The interface largely consists of two separate hydrophobic cores, one formed by the N-terminal helix bundles on top of the DNA binding groove, and the other by the β3–β5 sheet and α2–α3 below the DNA binding groove (Fig. 3). The upper core is formed by F41, V44, T76, and T83 from the left subunit (as viewed in Fig. 3B), and F52, L53, L56, and L57 from the right subunit.

The lower core is formed by F171, W216, and I218 from the left subunit, and I114, L118, and I126 from the right subunit (Fig. 3C). Both of these cores are surrounded by smaller sets of electrostatic interactions. At the upper rim, K40 and S77 of the left subunit form hydrogen bonds with D46 and N61 of the right subunit (Fig. 3B). At the lower rim, Q144 and R141 of the left subunit form hydrogen bonds with N127 and E135 of the right subunit (Fig. 3C). Most of the residues involved in the inter-subunit packing are conserved in distant homologs of LiRecT (Fig. S6), suggesting that the subunit packing and overall filament structure are likely to be conserved.

### Comparison to Rad52

The LiRecT structure permits a structure-based sequence alignment with Rad52 to identify the equivalent sets of residues used for interacting with DNA and neighboring subunits (Fig. S9). First and foremost, the inner strand in the complex with LiRecT closely overlays with the dT40 bound to the “inner” site of Rad52 (Fig. 2). Both strands are bound to the same position deep at the base of their respective grooves, where they are contacted by equivalent sets of residues extending from α3 (Figs. S9 and S10). Specifically, K206, R210, N211, and S214 from α3 of LiRecT correspond precisely to T148, K152, R153, and R156 from α3 of Rad52 (Figs. S10*C* and S10*F*). The outer strand of LiRecT approximately overlays with the ssDNA bound to the outer site of Rad52, but the latter is bound in a helical conformation that is clearly not poised for annealing (Figs. S10*B*, S10*D*, and S10*G*). Both proteins use the conserved β1–β2 hairpin to wedge into the DNA strands, and V98 from β1 of LiRecT is precisely equivalent to R55 from β1 of Rad52. In LiRecT the β1–β2 hairpin separates the base pairs by 9Å, whereas in Rad52 it separates the inner strand bases by 11Å (Figs. S10*E* and S10*H*). Although our structure of LiRecT captures the protein in a helical filament, and the structures of Rad52 reveal an 11-mer oligomeric ring, the two proteins use the same basic parts of their monomers for inter-subunit packing (Fig. S11), suggesting that the oligomers are related. At the sequence level, the most conserved part of the LiRecT and Rad52 structures is the interface between α2 and α3, which in both proteins is integral to the binding of the inner strand and the inter-subunit packing interactions. Finally, while the stoichiometry of the Rad52-ssDNA complex is 4 nt/monomer, the complex of LiRecT with annealed duplex has 5 bp/monomer. Whether the two proteins have slightly different stoichiometries, or there is a change in stoichiometry when the second strand is incorporated, remains to be determined.

### Structure of LiRecT in Complex with ssDNA

Prior work on λ-Redβ revealed that it binds ssDNA as oligomeric rings, and then forms helical filaments once a second complementary strand is added (16). To determine if there is a similar structural transition for LiRecT, we prepared a complex of it with just one 83-mer ssDNA and obtained a ~5Å resolution cryo-EM reconstruction by single particle analysis (Fig. S12). Surprisingly, the LiRecT-ssDNA complex also exists as lefthanded helical filaments, instead of as rings, but they are not as well ordered, and they do not stack end-to-end. Using a monomer of LiRecT from the complex with annealed duplex, 12 subunits of LiRecT could be docked into the reconstruction, although the density for the subunits at the ends of the filament is weaker (Fig. S12*C*). The filaments appear to be more tightly wound, with approximately 8 subunits per turn as opposed to ~10 for the complex with annealed duplex. Due to the lower resolution of this reconstruction, we could not fit the ssDNA to the map, although there is strong density for DNA in the groove above α2 and α3 (Fig. S12*D*). In addition, the density for the portion of each LiRecT subunit at the upper rim of the filament is particularly weak, for the entire length of the filament. This upper lobe of each monomer, which is formed by the αA–αC bundle and the β1–β2 hairpin, would likely clamp down on the DNA once the second strand is incorporated, to form the additional protein-DNA and inter-subunit interactions that are shown in Fig. 3 for the complex with annealed duplex. These interactions would further stabilize the filament complex to consolidate annealing. This provides a possible structural explanation for the dramatic increase in stability of the complex with two complementary strands that has been observed by gel-shift and single-molecule experiments for λ-Redβ (15,23).

### Analysis of LiRecT-DNA Complexes Formed in Solution

To determine if the complexes of LiRecT seen by cryo-EM also exist in solution, and in particular if the predicted full-length complex with two 83-mers is formed since the ends of the filament were not clear, mixtures of LiRecT protein alone, with 83-mer ssDNA, and with two complementary 83-mers added sequentially were analyzed by native mass spectrometry (nMS). Raw and deconvolved mass spectra for each sample are shown in Figs. S13 and S14, and a heat map summary of the oligomeric species formed for each protein-DNA mixture is shown in Fig. 4. Free LiRecT protein was largely monomeric at low concentration (1 μM), and while increasing the concentration to 30 μM resulted in some oligomer formation (up to 9-mers), no distinct oligomeric species was converged upon (Figs. 4 and S13). Mixing of LiRecT with one 83-mer ssDNA resulted in two types of complexes, one with 7-10 LiRecT subunits and one copy of the 83-mer (green in Fig. 4), and another with 15-17 subunits of LiRecT and two copies of the same 83-mer (blue in Fig. 4). Based on our previous results for λ-Redβ (17), we interpret the smaller complexes (green) as initial LiRecT-ssDNA substrate complexes, and the larger complexes (blue) as attempts at annealing at sites of partial complementarity. By contrast, mixing of LiRecT with the two complementary 83-mers added sequentially resulted in a more dominant complex containing 17 or 18 copies of LiRecT and one copy each of the 83+ and 83-oligonucleotides (purple in Fig. 4). Interestingly, the stoichiometry of the complex observed by nMS (83/17 or 83/18) is 4.9 or 4.6 bp/monomer, very close to the 5 bp/monomer observed for the cryo-EM structure. Moreover, complexes of LiRecT formed on slightly shorter pairs of complementary oligonucleotides (80- and 75-mers), contained 1-2 fewer subunits, as expected for a continuous oligomerization process like that of a helical filament. Interestingly, the complexes of LiRecT with just one ssDNA (green in Fig. 4) did not get noticeably smaller on the shorter ssDNAs, suggesting a different type of oligomerization process for the ssDNA complex.

**Fig. 4.**
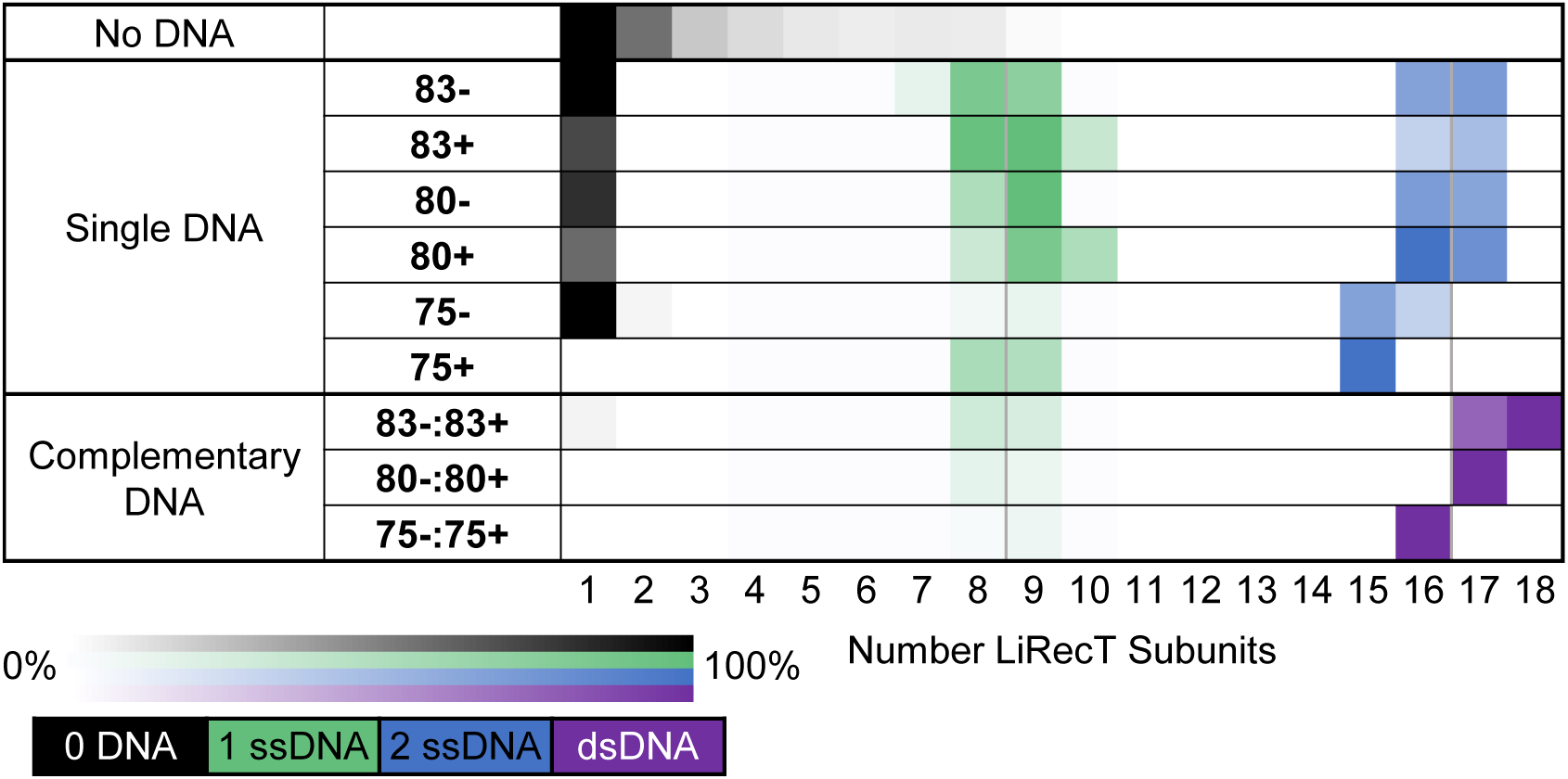
Native MS heat maps of LiRecT oligomers formed in solution. The first row (No DNA) shows the oligomers formed by 2 μM LiRecT in the absence of DNA. The second set of rows (Single DNA) shows the species formed after mixing a single DNA with 2 μM LiRecT. The third set of rows (Complementary DNA) shows the species formed after mixing two complementary DNAs sequentially with LiRecT. The order in which the DNAs are written corresponds to the order of addition. The heat maps indicate the relative intensities of all species present in each deconvolved spectrum (Fig. S13 and S14). The coloring corresponds to the DNA present in each complex: black to 0 ssDNA, green to 1 ssDNA, blue to two copies of the same ssDNA (2 ssDNA), and purple to one copy each of two complementary ssDNAs (dsDNA).

## Discussion

Using gel-shift assays with 33-mer and 83-mer oligonucleotides, Radding and colleagues discovered over 20 years ago that λ-Redβ exhibits unusual and intriguing DNA binding properties: it binds weakly to ssDNA, not at all to pre-formed dsDNA, but tightly to a duplex intermediate of annealing formed when two complementary oligonucleotides are added to the protein sequentially (15). They referred to this complex as an “intermediate” of annealing, rather than as a “product”, presumably because the DNA remained tightly bound to the protein. These experiments did not inform on the conformation of the bound DNA, and whether it was close to B-form, or adopted some other conformation, remained unknown. Our structure of LiRecT now reveals that the conformation is indeed quite distinct from B-form in being highly extended and underwound. Exactly where this conformation of DNA duplex lies along the energetic landscape of protein-mediated annealing (i.e. if it is a transition state or an intermediate), and whether or not it is a special conformation of DNA that is fundamental to annealing and common to all RecT/Redβ family members, remains to be determined.

Shortly after the unique DNA binding properties of λ-Redβ were discovered, oligomeric structures of λ-Redβ were visualized that closely paralleled the different DNA-bound states: rings for binding to ssDNA, and helical filaments for binding to annealed duplex (16). The filaments of LiRecT that we have observed by cryo-EM closely match the filaments of λ-Redβ seen by negative stain EM: they are left-handed, and have similar dimensions and helical parameters. Given that LiRecT and λ-Redβ share limited sequence identity with one another (<15%), the fact that they share a conserved helical filament structure would tend to suggest that the conformation of the duplex intermediate that is bound to them is also conserved.

Egelman and colleagues predicted that the duplex intermediate formed by λ-Redβ was likely to be bound along the inner surface of the helical filament (though not along the helical axis), based on the observation that it was protected from DNAse I cleavage (15,16). Our structure of LiRecT reveals that it is instead bound to a groove that runs along the outer surface of the filament. The fact that the DNA is buried in such a deep and narrow groove, and that its conformation is far from B-form, may explain why it is still protected from DNAse I cleavage.

We have so far not been able to visualize oligomeric rings of LiRecT bound to ssDNA, like the 11-mer rings seen for λ-Redβ (16) and Rad52 (10–12). Our nMS data indicate that LiRecT exists in a monomer-oligomer equilibrium (up to 9-mer) in the absence of DNA, and as a complex of 7 to 10 subunits on a single 83-mer ssDNA. Interestingly, in the complexes with a single 83-mer ssDNA, LiRecT does not appear to bind to the full-length of the DNA, as it does for the complex with annealed duplex. These observations are similar to our previous nMS analysis of λ-Redβ (17), although the latter protein had a higher propensity to form oligomers in the absence of DNA. We favor a model in which RecT/Redβ proteins oligomerize weakly and dynamically on their own, assemble onto ssDNA as clusters of cooperatively bound monomers to form partial rings or filaments, and as stable helical filaments once the complementary strand is incorporated into the complex. The weaker complexes on ssDNA may allow for dynamic sampling with additional strands of ssDNA until a complementary sequence is found and aligned, at which point the N-terminal lobe of each protein monomer clamps down on the DNA to stabilize the complex and consolidate annealing.

Filaments of both λ-Redβ and LiRecT can be several helical turns in length, but annealing assays with λ-Redβ indicate that the minimal length needed for successful annealing *in vitro* is only 20 bp (17,18). Moreover, oligonucleotides as short as 70-mers are fully functional for Redβ annealing *in vivo*. Therefore, we consider it unlikely that long helical filaments of these proteins would form *in vivo*. Although the helical filament is highly stable, at least as compared to the ssDNA complexes (15,23), it would likely disassemble *in vivo* once the two DNA molecules are spliced together, possibly due to the greater torsional stress of being bound to the middle of a larger DNA duplex, as opposed to at the ends. An alternative possibility is that a DNA helicase, or a component of the DNA replication machinery could be involved in removing the protein from the DNA *in vivo*.

Residues 1-33 and 225-271 of LiRecT were not resolved in our 3Dreconstruction of the filament. These residues are however part of a RoseTTAFold model for the LiRecT monomer, as shown in magenta in Figure *S4B*. Residues 1-33 form two α-helices, one that is quite long (residues 1-27) and extends away from the core of the monomer, and another that is short (residues 27-33) and forms a right angle with αA. In our reconstruction, there is density for what appears to be a helix preceding αA. Although the density for this helix was not clear enough to model, it appears to pack against αA of the neighboring subunit, and thereby add to the inter-subunit contacts. There is no sign of density that would correspond to the long N-terminal α-helix from the RoseTTAFold model.

By analogy with λ-Redβ, it is likely that the extra residues at the C-terminal end (225-271) fold into a small helical domain for forming interactions with partner proteins, including the host single-stranded DNA-binding protein (SSB) (24). In the RoseTTAFold model, residues 242-271 extend away from the filament to possibly form such a domain, but residues 220-238 form an α-helix that packs against the β1–β2 hairpin and splits right into the DNA strands. The placement of this helix is not consistent with DNA binding, but it could conceivably adopt this position in the LiRecT monomers before they assemble onto the DNA. Further studies will be needed to resolve these issues.

The structure confirms that the RecT/Redβ family of annealing proteins is indeed homologous to Rad52. The two proteins bind to the first ssDNA in a highly conserved manner, with equivalent sets of residues contacting the DNA from conserved secondary structural elements (α2 and α3). Moreover, the proteins use approximately the same portions of their monomers for inter-subunit packing, suggesting that their oligomers are related. However, Rad52 has so far only been observed to form rings, and has not been seen to form helical filaments. Rad52 also exhibits somewhat different DNA-binding properties from λ-Redβ in binding with higher affinity to ssDNA and to pre-formed dsDNA (13). Furthermore, a distinct complex of Rad52 with a duplex intermediate of annealing like those of λ-Redβ and LiRecT has not yet been observed. Nonetheless, the DNA binding grooves on the LiRecT and Rad52 structures are formed by the same set of secondary structural elements, and are similarly deep and narrow, suggesting that a complex of Rad52 with two strands of complementary DNA bound simultaneously could very well be formed. The existence of such a complex would favor a *cis* mechanism of annealing in which the two DNA strands are bound to the same protein oligomer as they are annealed to one another, as opposed to a *trans* mechanism in which annealing is mediated by interaction of two separate Rad52-ssDNA complexes.

Although human Rad52 has been widely considered to exist as stable oligomeric rings, yeast Rad52 is expressed at only nanomolar concentrations *in vivo* (25), and human Rad52 is largely monomeric at sub-micromolar concentrations *in vitro* (26). Thus, non-ring forms of Rad52 could still be relevant to its mechanism.

Some features of the LiRecT-DNA complex are remarkably similar to other types of DNA recombination proteins. The 1.5x extended conformation of DNA and the 5 bp repeating pattern of extension are similar to the triplet-repeating conformation of DNA bound to *E. coli* RecA protein (27). The LiRecT-DNA complex also shares some remarkably similar features with a multi-subunit complex of *E. coli* Cascade bound to an RNA-DNA duplex hybrid (28). In Cascade, the duplex is bound to a very similar groove along the outer surface of a right-handed helical assembly of subunits. The duplex is similarly extended and under-wound, bound in a pattern that repeats every 6 bp steps due to a similar β-hairpin insertion, and has the first strand added (RNA) in the deepest part of the groove, and the second strand added (DNA) at the outer part of the groove. These striking similarities of LiRecT with functionally (but not structurally) related proteins point to fundamental principles of DNA transactions that are still being unraveled.

## Materials and Methods

### Protein expression and purification

The gene expressing LiRecT (UniProtKB – Q92FL9) was PCR amplified from *Listeria innocua* CLIP 11262 genomic DNA (ATCC BAA-680) and cloned into pET28b between the *Nde*I and *BamH*I restriction sites to express a protein with an N-terminal 6His-tag and a site for thrombin cleavage. The protein was expressed in BL21(AI) *E. coli* cells (Invitrogen) in 6 × 1L cultures at 37°C grown to an optical density at 600 nm of 0.65, and induced by 1 mM IPTG and 0.2% arabinose. At four hours post-induction, the cells were harvested by centrifugation, resuspended in 60 ml of Buffer A (50 mM NaH_2_PO_4_, 300 mM NaCl, 10 mM imidazole, pH 8.0) and frozen at −80°C. After thawing, lysozyme (1 mg/ml), PMSF (0.1 mg/ml), leupeptin and pepstatin (1 μg/ml each) were added and incubated for 60 min on ice. The cells were then sonicated on ice, centrifuged at 38,000 × g for 3 × 30 min, and the final supernatant was loaded on to a 2 × 5 ml HisTrap column (Cytiva) at 0.5 ml/min. The column was washed with 30 ml of Buffer A, 200 ml of Buffer A containing 30 mM imidazole, and eluted with a 200 ml gradient of 30-500 mM imidazole in Buffer A. After SDS-PAGE analysis, pooled fractions were mixed with 100 units of Thrombin (Cytiva), dialyzed at room temperature into Buffer B (20 mM NaH_2_PO_4_, 1500 mM NaCl, pH 7.4), and loaded back onto the HisTrap column. The flow through was collected, dialyzed at 4°C into Buffer C (20 mM Tris pH 8.0) and loaded onto a 2 × 5 ml QFF column (Cytiva) at 1 ml/min. After washing with Buffer C for 30 ml, the protein was eluted with a 100 ml gradient to Buffer C plus 1M NaCl. Pooled fractions were dialyzed into Buffer D (20 mM Tris, 1 mM DTT, pH 8.0), concentrated to 50 mg/ml (Vivaspin 20, 10 kDa MWCO), and stored at −80°C in 50 μl aliquots. Protein concentration was determined by O.D. at 280 nm using an extinction coefficient of 43,890 M^−1^ cm^−1^, which was determined from the amino acid sequence, which has 5 tryptophan residues.

### DNA annealing assay

A gel shift assay with two complementary 50-mer oligonucleotides labeled at the 5’-end with either Cy3 or Cy5 was performed as described previously for λ-Redβ (17). All oligonucleotides used in this study were purchased HPLC-purified from Integrated DNA Technologies, dissolved in ddH_2_O, and stored at −20°C. Their full sequences are as described previously (17). Briefly, 10 μM of Redβ in PBS was mixed with 50 μM (nt) of the indicated oligonucleotide and incubated at 37°C for 30 min. For some samples, as indicated on the gel (lanes labeled “ad” or “nc”), a second oligonucleotide was added, and incubated for an additional 30 min at 37°C. For all samples, the total reaction volume was 30 μl. For visualization, 3 μl of each complex was mixed with 7 μl Orange G dye (65% w/v sucrose, 10 mM Tris-HCl pH 7.5, 10 mM EDTA, 0.3% Orange G powder from Sigma Life Sciences), loaded onto a 1.0% agarose gel, and electrophoresed in 2x TBE for 40 min at 90 V. Gels were imaged using an Sapphire Biomolecular Imager (Azure Biosystems).

### Cryo-EM Sample Preparation

The complex of LiRecT with annealed duplex was prepared by first incubating 0.7 mg/ml protein with one 83-mer oligonucleotide (83-) at a ratio of 4 nt/monomer (94 μM nucleotides) in 20 mM KH_2_PO_4_, 10 mM MgCl_2_ pH 6.0 at 37 °C for 15 min. Then an equivalent amount of the complementary 83-mer (83+) was added and incubated for an additional 15 min at 37°C, after which the prepared complex was kept on ice for approximately 90 min. The complex of LiRecT with ssDNA was prepared in the same manner as that for annealed duplex, but only the first strand of ssDNA (83-) was added. For both complexes the total reaction volume was 19 μl. Just prior to vitrification, 1 μl of 1.5 mM n-dodecyl-beta-maltoside (Anatrace; final concentration at 0.5 CMC) was added and incubated for 30 seconds, and then 4 μl of the mixture was added to a Quantifoil R1.2/1.3 Au 300 mesh grid (Electron Microscopy Sciences) that had been glow discharged for 60 seconds at 20 mA using a Pelco easiGlow. After applying the sample, the grid was immediately frozen by plunging into liquid ethane using a Vitrobot Mark IV (Thermo Fisher Scientific) at 4 °C, 100% humidity, 1.5 second blot time, and 0 blot force. Ted Pella 595 filter paper (Product # 47000-100) was used for blotting.

### Cryo-EM Data Acquisition

For the complex with 83-mer annealed duplex, images were collected on a 300 keV Titan Krios G3i electron microscrope (Thermo Fisher Scientific) operating in nanoprobe EFTEM mode with 50 μm C2 aperture, 100 μm objective aperture, a Gatan BioContinuum energy filter (20 eV slit width, zero energy loss), a Cs corrector, and a Gatan K3 direct electron detector operating in counting mode. Automated data collection was performed in EPU with defocus values ranging from −1 to −3.5 μm at a magnification of 81,000x and a pixel size of 0.899 Å (non-super-resolution). The dose rate was adjusted to 24.28 e-/Å^2^ /s with an exposure time of 2.7 s split into 36 fractions to achieve a total dose of 66 e-/Å^2^. A total of 2038 movies were collected. For the complex with 83-ssDNA, the same settings were used, except for the following: the data were collected in super-resolution mode such that the pixel size was 0.4495 Å, the does rate was adjusted to 22.80 e-/Å^2^/s with an exposure time of 2.83 s split into 45 fractions for a total dose of 65 e-/Å^2^, and 1619 movies were collected.

### Cryo-EM Data Processing

For the data for the complex with annealed duplex, movies were imported into cryoSPARC v2.15.0 (29) for single particle analysis. Patch motion correction was implemented with a 3Å maximum alignment resolution and a B-factor of 500. Patch CTF estimation was implemented with an amplitude contrast of 0.1. From the motion- and CTF-corrected micrographs, approximately 1000 particles were manually picked and used for one round of 2D classification. Six 2D class averages representing different particle orientations were chosen and used as templates for automated particle picking, which resulted in approximately 1,100,000 particles. Particles were extracted with a box size of 210 pixels and put through three rounds of 2D classification to result in 391,275 cleaned particles. The cleaned particles were used to generate an initial model with *ab-initio* reconstruction, which was refined in homogenous refinement to yield a 3D reconstruction with an FSC gold standard resolution of 3.4Å (tight mask), or 4.3Å (no mask). After polishing, the resulting 3D reconstruction showed clear density for protein backbone, side chains, and two strands of DNA including bases. The data for LiRecT with ssDNA were processed in the same manner to result in 180,965 cleaned particles, and a final resolution of 4.5Å (tight mask). This resolution is likely over-estimated however, as the FSC curve was oscillating.

### Model Building and Refinement

For the complex with annealed duplex, the two unmasked half maps from cryoSPARC were input into the RESOLVE procedure for density modification in PHENIX version 1.20.1-4487 (30) which improved the resolution by 0.22Å from 3.81Å (FSCref=0.5) to 3.59Å (FSCref=0.5). A model of one protein monomer containing residues 34-224 (out of 271 total) was built into the central portion of the filament with COOT version 0.8.7 (31), and then transformed iteratively into density for nine neighboring subunits using CHIMERA version 1.13.1 (32). Additional subunits towards the ends of the filament were visible in the reconstruction, but not included in the final model, as the density for these regions was progressively weaker. The 3D reconstruction also showed clear density for 48 bp of DNA duplex at the central portion of the filament, which was also built using COOT. Once a 10-subunit filament was built, the NCS operators were determined from the structure using Find NCS in PHENIX, and then used for 10-fold NCS averaging in Resolve, which further increased the resolution to 3.50 Å (FSCref=0.5). The final model consists of 10 protein subunits and 48 bp of DNA. Real space refinement and model validation in PHENIX yielded a final FSC=0.143 map to model resolution of 3.2 Å. During refinement, 10-fold NCS constraints were applied to the protein monomers, but not to the DNA. Final refinement and model validation statistics are shown in Table S1. For the structure with 83-mer ssDNA, the resolution of the reconstruction did not enable the model to be built from scratch as secondary structures were barely visible, but six LiRecT subunits from the structure with annealed duplex could be auto-fit into density using CHIMERA, and additional subunits could be fit using PHENIX (dock_in_map). The final model consisting of 8 LiRecT subunits was refined in PHENIX by rigid body refinement only. Structural figures were prepared using PyMOL version 2.5 (33). Atomic coordinates and maps have been deposited in EMDB under accession codes EMD-26434 (complex with 83-mer annealed duplex) and EMD-26437 (complex with 83-mer ssDNA).

### Native Mass Spectrometry

LiRecT protein was buffer exchanged into 100 mM ammonium acetate pH 7 (unadjusted) using Micro BioSpin P6 spin columns (Bio-Rad Laboratories, Hercules, CA, USA). All ssDNAs were dialyzed into 100 mM ammonium acetate with Pierce 96-well microdialysis devices with 3.5K MWCO (Thermo Fisher Scientific). For preparation of LiRecT-DNA complexes, LiRecT was diluted to the experimental concentrations indicated, and then the first ssDNA was added at the indicated concentration based on nucleotides (nt) per monomer of LiRecT, and incubated at 37°C for at least 15 min. For complexes with annealed duplex, the second complementary ssDNA was then added and incubated for an additional 15 min. Samples (3 – 5 μl) were directly loaded into nanoESI emitters that were pulled in-house from borosilicate filament capillaries (OD 1.0 mm, ID 0.78 mm, Sutter Instrument) using a P-97 Flaming/Brown Micropipette Puller (Sutter Instrument). Experiments were performed on a Thermo Scientific Q Exactive Ultra-High Mass Range (UHMR) mass spectrometer from Thermo Fisher that was modified to allow for surface-induced dissociation (SID, not used in this work) similar to a previously described modification (34). The same instrument settings were used as described previously (34). Ion activation was necessary for improved transmission and de-adducting of ions to resolve species at higher m/z. For this, in-source trapping (IST) of −10 V and higher energy collisional dissociation (HCD) of 90 V was used for the LiRecT plus DNA mixtures. All data were deconvolved using UniDec V4.4 (35). A range of deconvolution settings was initially surveyed. The settings optimized for LiRecT plus DNA mixtures were the following: 2000 to 16000 m/z, charge range of 1 to 70, mass range of 10 to 800 kDa, sample mass every 10 Da, split Gaussian/Lorentzian, peak FWHM 3 or 4 Th, artifact suppression 40, charge smooth width 2.0, point smooth width 2, and native charge offset −20 to 10 or 20. The use of manual mode to assign a fraction of the peaks with charge states was needed to reduce artifacts. The resulting deconvolutions were plotted as relative signal intensities.

## Acknowledgments

This work was funded by grants from the National Science Foundation (MCB-1616105 to C.E.B) and National Institutes of Health (T32GM11829 to B.J.C. and P41GM128577 to V.H.W). The content is solely the responsibility of the authors and does not necessarily represent the official views of the National Science Foundation or the National Institutes of Health.

