## Supplementary Information Appendix for "Structure of a Rad52 homolog from bacteriophage in complex with a novel duplex intermediate of DNA annealing"

**This PDF file includes:**

Figures S1 to S14
Table S1
Legend for Movie S1
SI References

**Other supplementary materials for this manuscript include the following:**

Movie S1

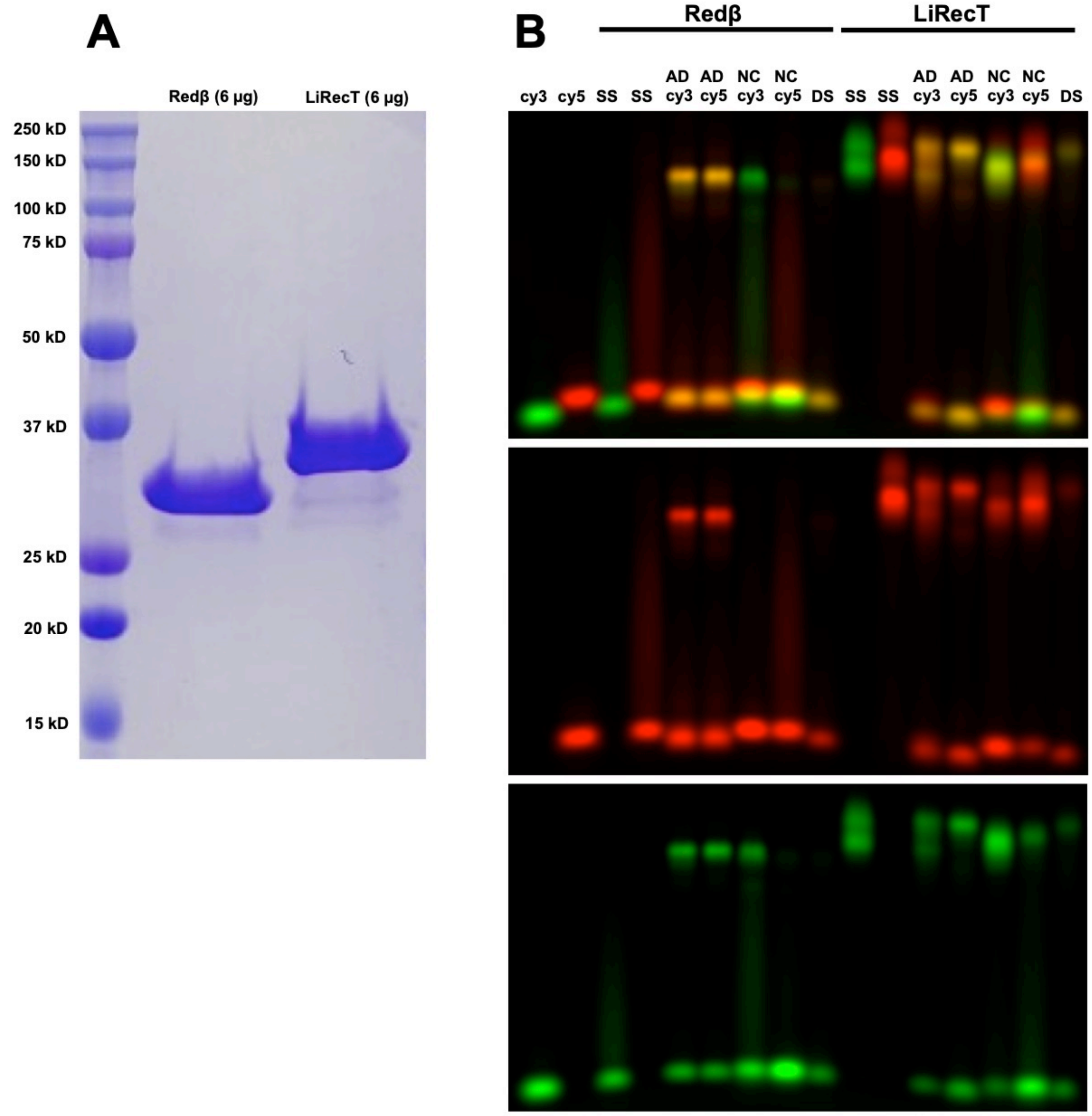


**Fig. S1**. **Purification and DNA binding properties of LiRecT**. (***A***) 12.5% SDS-PAGE gel stained with Coomassie Blue R250. (***B***) Gel-based DNA annealing assay comparing the DNA binding activities of LiRecT with λ-Redβ. Lanes contain the following complexes: cy3 and cy5: single 5'-labeled 50-mer oligo at 50 µM (nt) in the absence of protein; SS: a single oligo at 50 µM with 10 µM protein; AD cy3 and AD cy5: sequential addition of two complementary oligos, each at 50 µM, to 10 µM protein (with the indicated oligo added first); NC cy3 and NC cy5, sequential addition of two non-complementary oligos to protein; DS: addition of two complementary pre-annealed oligos to protein. The middle and lower panels are the single channel exposures, to show the oligos present (Cy3-green and/or Cy5-red) in each species. Notice that λ-Redβ binds weakly to ssDNA (as evidenced by the streaking in the SS lanes), forms a distinct complex containing both oligos (yellow band) when two complementary oligos are added to the protein sequentially (AD), does not form this complex when the two oligos are non-complementary (NC), and forms no interaction with pre-formed dsDNA (DS). LiRecT exhibits similar behavior as λ-Redβ, but binds more tightly to ssDNA. Sequences of all oligonucleotides are as described previously (1).

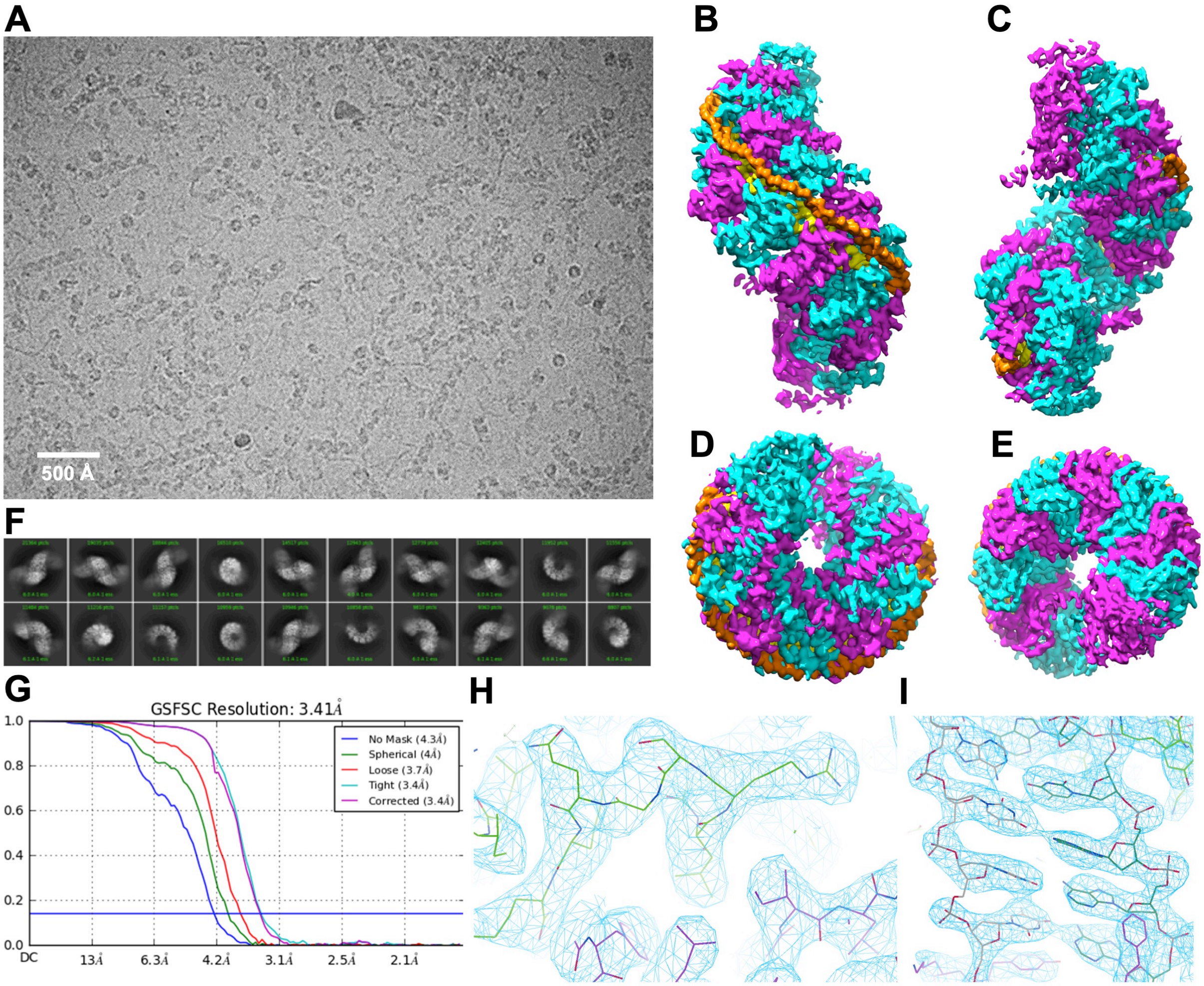


**Fig. S2: CryoEM structure determination of LiRecT bound to 83-mer annealed duplex.** (***A***) Example Krios-K3 image showing side and top views of the LiRecT filament. (***B-E***) 3D reconstruction of the filament showing 10 subunits of LiRecT alternating in magenta and cyan. The inner strand of DNA is in yellow and the outer strand is in orange. The four panels show views from the front (B), back (C), top (D) and bottom (E). (***F***) Selected 2D class averages from 390,000 cleaned particles (***G***) FSC plot after homogenous refinement in cryoSPARC. (***H, I***) Cose-up views of density for representative regions of protein and DNA, respectively. Notice that the density clearly allows for placement of side chains and bases, although the sequence of the DNA could not be interpreted, presumably due to difficulty in alignment of particles along the filament axis. Figures in panels B-E were generated by Chimera (2). Figures in panels F and G were generated by cryoSPARC (3). Figures in panels H and I were generated by Coot (4).

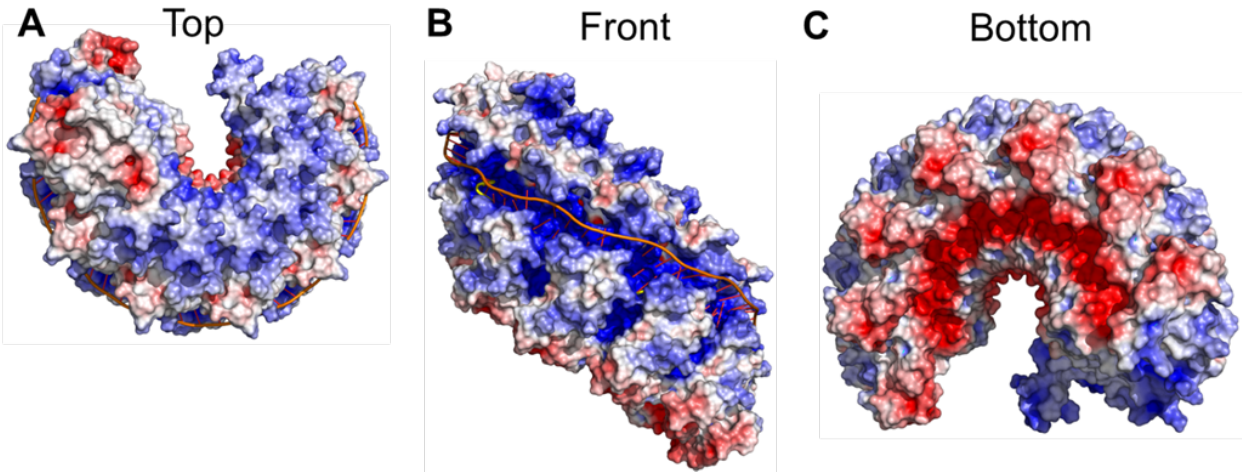


**Fig. S3. Electrostatic surface views of the LiRecT filament.** Colors of the electrostatic surface correspond to regions of positive (blue), neutral (white), and negative (red) charge. The upper surface of the filament is mostly positively charged (***A***), the outer surface contains a deep and narrow positively charged groove (***B***), and the bottom surface (***C***) is mostly negatively charged. The complementary charges could help to promote the end-to-end stacking of the filaments that is observed in the EM images. Figures were generated with PyMOL (5).

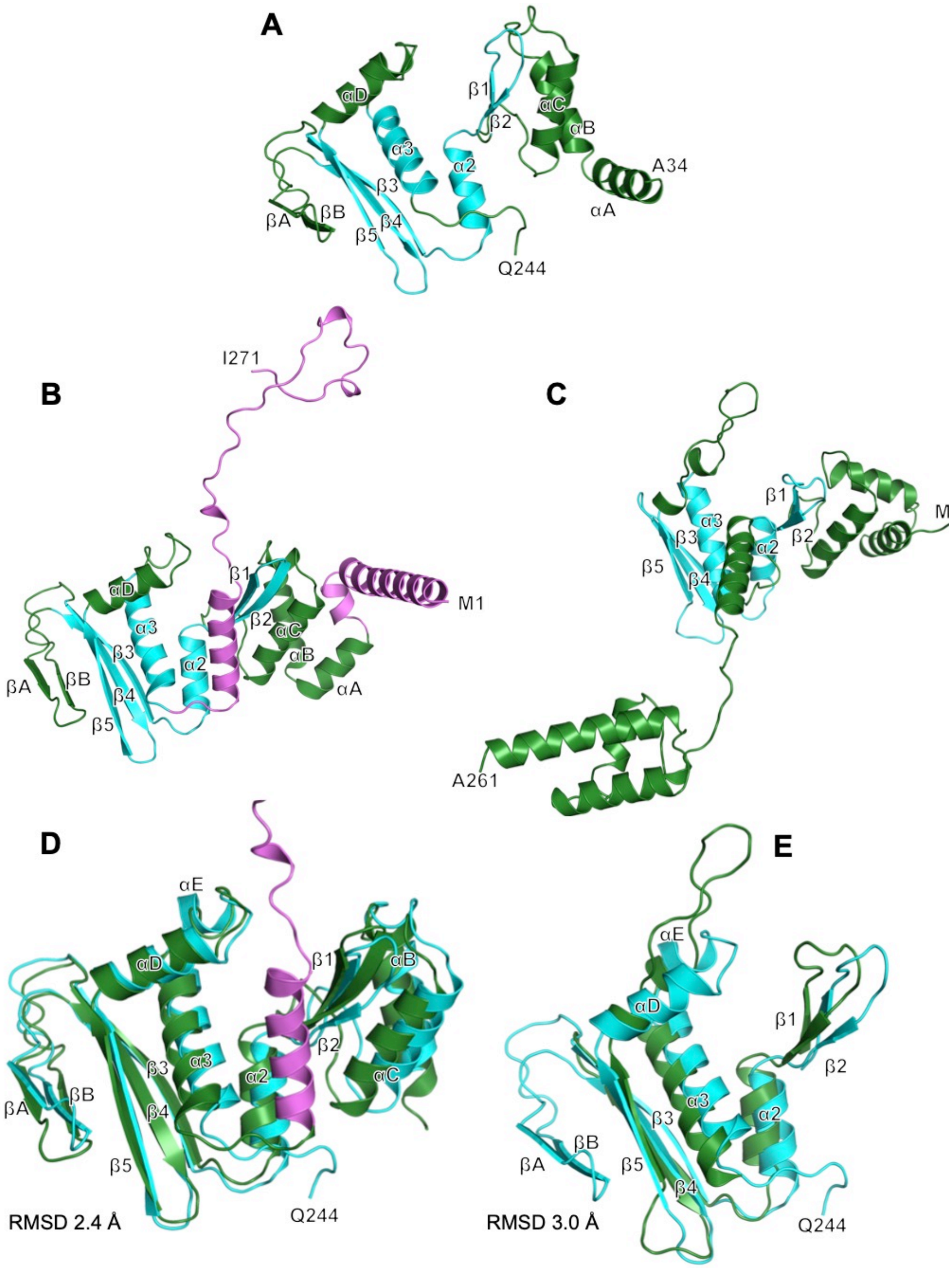


**Fig. S4**. **Comparison of the LiRecT cryo-EM structure with the structures of LiRecT and λ-Redβ predicted by RoseTTAFold**. (***A***) Ribbon diagram of the LiRecT monomer from the cryo-EM structure. The core that is common to Rad52 is colored in cyan. (***B***) Structure of LiRecT predicted by RoseTTAFold (6). Extra segments predicted by RoseTTAFold that are absent from the cryo-EM structure are colored magenta. (***C***) Structure of λ-Redβ predicted by RoseTTAFold. The helical bundle at the lower left is the C-terminal domain. (***D***) Alignment of the cryo-EM (cyan) and RoseTTAFold (green/magenta) structures of LiRecT. (***E***) Alignment of the cores of the LiRecT cryo-EM (cyan) and λ-Redβ RoseTTAFold (green) structures. Notice that λ-Redβ is missing the βA–βB insertion of LiRecT and has an extended loop between β5 and α3 (in place of αD and αE).

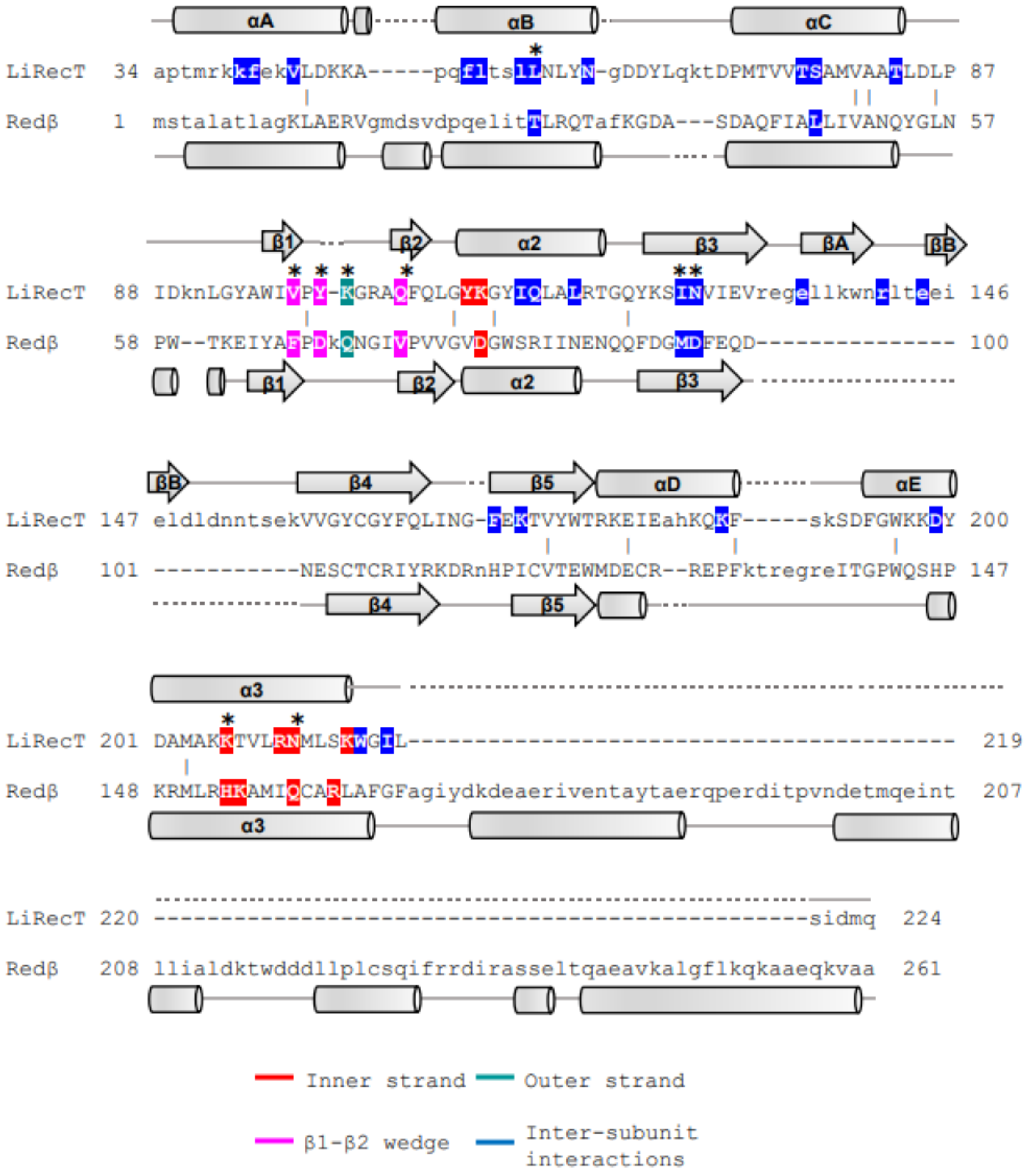


**Fig. S5. Structure-based sequence alignment of LiRecT with a model of λ-Redβ predicted by RoseTTAFold.** Structurally equivalent residues are in uppercase, structurally non-equivalent residues (including insertions) are in lowercase. Amino acid identities are indicated by vertical bars. The shading indicates amino acids of LiRecT that contact the inner strand (red), the outer strand (teal), the DNA from the β1-β2 hairpin insertion (violet), and form inter-subunit interactions (blue). The ∗ symbols indicate key functional residues of LiRecT and their predicted structurally conserved counterparts in λ-Redβ.

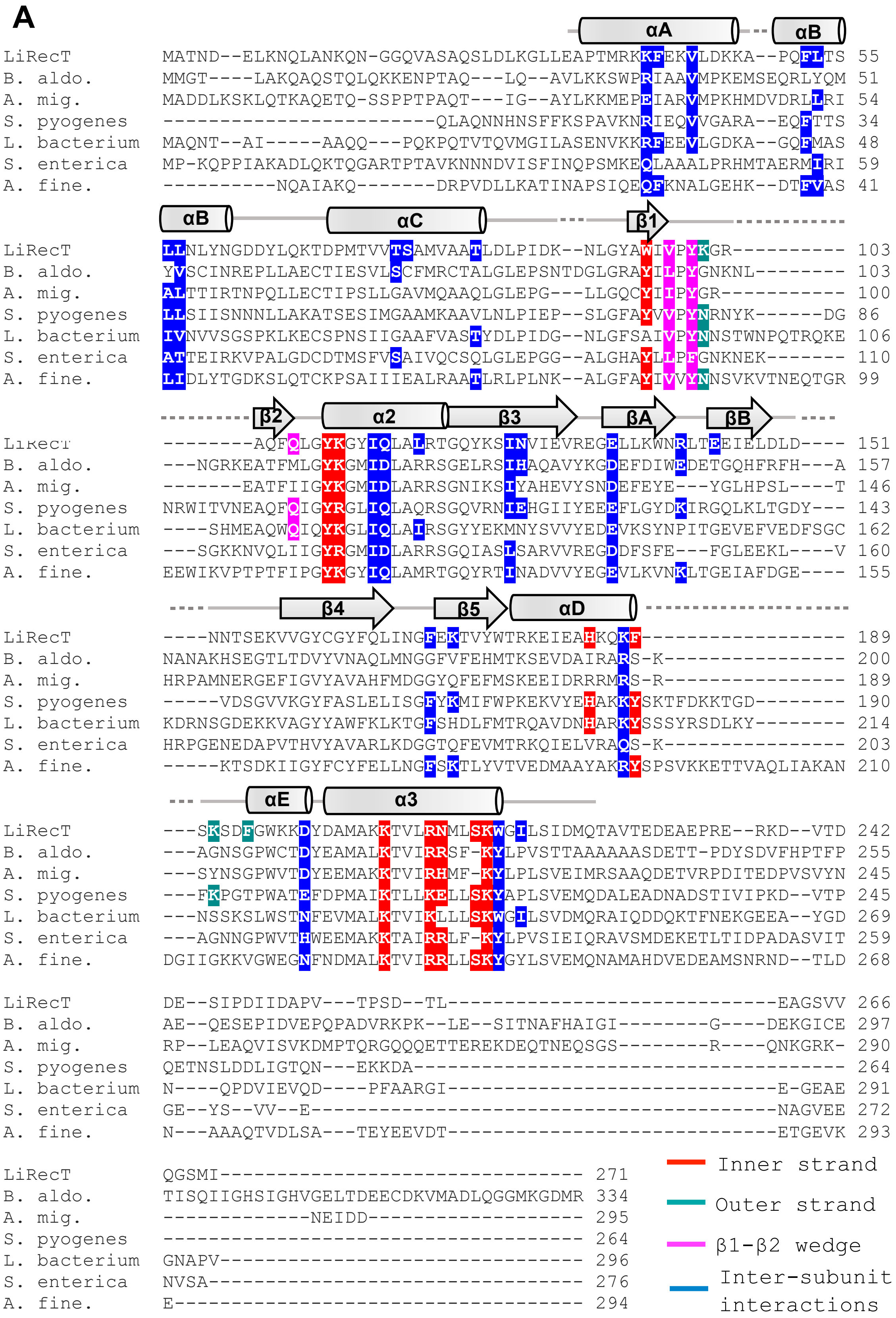

A
αA
αB
LiRecT MATND--ELKNQLANKQN-GGQVASAQSLDLKGLLEAPTMRKKFEKVLDKKA--PQFLTS 55
B. aldo. MMGT----LAKQAQSTQLQKKENPTAQ---LQ--AVLKKSWPRIAAVMPKEMSEQRLYQM 51
A. mig. MADDLKSKLQTKAQETQ-SSPPTPAQT---IG--AYLKKMEPEIARVMPKHMDVDRLLRI 54
S. pyogenes -------------------------QLAQNNHNSFFKSPAVKNRIEQVVGARA--EQFTTS 34
L. bacterium MAQNT--AI-----AAQQ---PQKPQTVTQVMGILASENVKKRFEEVLGDKA--GQFMAS 48
S. enterica MP-KQPPIAKADLQKTQGARTPTAVKNNNDVISFINQPSMKEQLAAALPRHMTAERMIRI 59
A. fine. ----------NQAIAKQ-------DRPVDLLKATINAPSIQEQFKNALGEHK--DTFVAS 41
αB
αC
β1
LiRecT LLNLYNGDDYLQKTDPMTVVTSAMVAATLDLPIDK--NLGYAWIVPYKGR---------- 103
B. aldo. YVSCINREPLLAECTIESVLSCFMRCTALGLEPSNTDGLGRAYILPYGNKNL-------- 103
A. mig. ALTTIRTNPQLLECTIPSLLGAVMQAAQLGLEPG---LLGQCYIIPYGR----------- 100
S. pyogenes LLSIISNNNLLAKATSESIMGAAMKAAVLNLPIEP--SLGFAYVVPYNRNYK------DG 86
L. bacterium IVNVVSGSPKLKECSPNSIIGAAFVASTYDLPIDG--NLGFSAIVPYNNSTWNPQTRQKE 106
S. enterica ATTEIRKVPALGDCDTMSFVSAIVQCSQLGLEPGG--ALGHAYLIPFGNKNEK------- 110
A. fine. LIDLYTGDKSLQTCKPSAIIIEALRAATLRLPLNK--ALGFAYIVVYNNSVKVTNEQTGR 99
β2
α2
β3
βA
βB
LiRecT --------AQFQLGYKGYIQLALRTGQYKSINVIEVREGELLKWNRLTEEIELDLD---- 151
B. aldo. ---NGRKEATFMLGYKGMIDLARRSGELRSIHAQAVYKGDEFDIWEDETGQHFRFH---A 157
A. mig. -------EATFIIGYKGMIDLARRSGNIKSIYAHEVYSNDEFEYE---YGLHPSL----T 146
S. pyogenes NRWITVNEAQFQIGYRGLIQLAQRSGQVRNIEHGIIYEEEFLGYDKIRGQLKLTGDY--- 143
L. bacterium ----SHMEAQWQIQYKGLIQLAIRSGYYEKMNYSVVYEDEVKSYNPITGEVEFVEDFSGC 162
S. enterica ---SGKKNVQLIIGYRGMIDLARRSGQIASLSARVVREGDDFSFE---FGLEEKL----V 160
A. fine. EEWIKVPTPTFIPGYKGYIQLAMRTGQYRTINADVVYEGEVLKVNKLTGEIAFDGE---- 155
β4
β5
αD
LiRecT ----NNTSEKVVGYCGYFQLINGFEKTVYWTRKEIEAHKQKF------------------ 189
B. aldo. NANAKHSEGTLTDVYVNAQLMNGGFVFEHMTKSEVDAIRARS-K---------------- 200
A. mig. HRPAMNERGEFIGVYAVAHFMDGGYQFEFMSKEEIDRRRMRS-R---------------- 189
S. pyogenes -----VDSGVVKGYFASLELISGFYKMIFWPKEKVYEHAKKYSKTFDKKTGD-------- 190
L. bacterium KDRNSGDEKKVAGYYAWFKLKTGFSHDLFMTRQAVDNHARKYSSSYRSDLKY-------- 214
S. enterica HRPGENEDAPVTHVYAVARLKDGGTQFEVMTRKQIELVRAQS-K---------------- 203
A. fine. -----KTSDKIIGYFCYFELLNGFSKTLYVTVEDMAAYAKRYSPSVKKETTVAQLIAKAN 210
αE
α3
LiRecT ---SKSDFGWKKDYDAMAKKTVLRNMLSKWGILSIDMQTAVTEDEAEPRE--RKD--VTD 242
B. aldo. ---AGNSGPWCTDYEAMALKTVIRRSF-KYLPVSTTAAAAAASDETT-PDYSDVFHPTFP 255
A. mig. ---SYNSGPWVTDYEEMAKKTVIRHMF-KYLPLSVEIMRSAAQDETVRPDITEDPVSVYN 245
S. pyogenes ---FKPGTPWATEFDPMAIKTLLKELLSKYAPLSVEMQDALEADNADSTIVIPKD--VTP 245
L. bacterium ---NSSKSLWSTNFEVMALKTVIKLLLSKWGILSVDMQRAIQDDQKTFNEKGEEA--YGD 269
S. enterica ---AGNNGPWVTHWEEMAKKTAIRRLF-KYLPVSIEIQRAVSMDEKETLTIDPADASVIT 259
A. fine. DGIIGKKVGWEGNFNDMALKTVIRRLLSKYGYLSVEMQNAMAHDVEDEAMSNRND--TLD 268
LiRecT DE--SIPDIIDAPV---TPSD--TL-----------------------------EAGSVV 266
B. aldo. AE--QESEPIDVEPQPADVRKPK--LE--SITNAFHAIGI--------G----DEKGICE 297
A. mig. RP--LEAQVISVKDMPTQRGQQQETTEREKDEQTNEQSGS--------R----QNKGRK- 290
S. pyogenes QETNSLDDLIGTQN---EKKDA---------------------------------------- 264
L. bacterium N---QPDVIEVQD----PFAARGI-----------------------------E-GEAE 291
S. enterica GE--YS--VV--E-----------------------------------------NAGVEE 272
A. fine. N---AAAQTVDLSA---TEYEEVDT-----------------------------ETGEVK 293
LiRecT QGSMI------------------------------- 271
B. aldo. TISQIIGHSIGHVGELTDEECDKVMADLQGGMKGDMR 334
A. mig. -------------NEIDD------------------- 295
S. pyogenes ------------------------------------ 264
L. bacterium GNAPV------------------------------- 296
S. enterica NVSA-------------------------------- 276
A. fine. E----------------------------------- 294
Inner strand
Outer strand
β1-β2 wedge
Inter-subunit interactions

**B**

| | Li | Ba | Am | Sp | Lb | Se | Af |
|---|---|---|---|---|---|---|---|
| **LiRecT** | 100 | 25.38 | 24.60 | 36.02 | 37.69 | 26.48 | 41.83 |
| **B. aldo.** | 25.38 | 100 | 44.06 | 23.75 | 22.26 | 25.38 | 24.02 |
| **A. mig.** | 24.60 | 44.06 | 100 | 27.51 | 25.40 | 24.60 | 23.05 |
| **S. pyogenes** | 36.02 | 23.75 | 27.51 | 100 | 34.90 | 36.02 | 31.54 |
| **L. bacterium** | 37.69 | 22.26 | 25.40 | 34.90 | 100 | 37.69 | 34.19 |
| **S. Enterica** | 26.48 | 36.60 | 43.75 | 24.57 | 23.64 | 100 | 23.58 |
| **A. fine.** | 41.83 | 24.02 | 23.05 | 31.54 | 34.19 | 41.83 | 100 |

**Fig. S6. Position specific iterative blast alignment of seven LiRecT homologs. (*A*)** Multiple sequence alignment from Clustal-Omega (7). The homologs were chosen on the basis of having low to moderate levels of sequence identity with LiRecT and with one another (see panel *B*). Secondary structure elements of LiRecT are shown above the alignment. The shading indicates amino acids in LiRecT that contact the inner strand (red), the outer strand (teal), form the β1-β2 wedge (violet), and form inter-subunit interactions (blue). These residues are also shaded in the sequence of the homologs that contain a similar amino acid type at that position. (***B***) Matrix showing the pair-wise sequence identities for all seven homologs. All homologs share from 23-44% pairwise sequence identity with LiRecT and with one another. The abbreviations to the left of each sequence correspond to: B. Aldo., *Bifidobacterium adolescentis* (WP_147538225.1); A. mig., *Aneurinibacillus migulanus* (WP_043070608.1); S. pyogenes, *Streptococcus pyogenes* (WP_115222948.1); L. bacterium, *Lachnospiraceae bacterium* (MBQ9437646.1); S. enterica, *Salmonella enterica* (EAO8776355.1); A. fine., *Alistipes finegoldii* (WP_014774661.1)

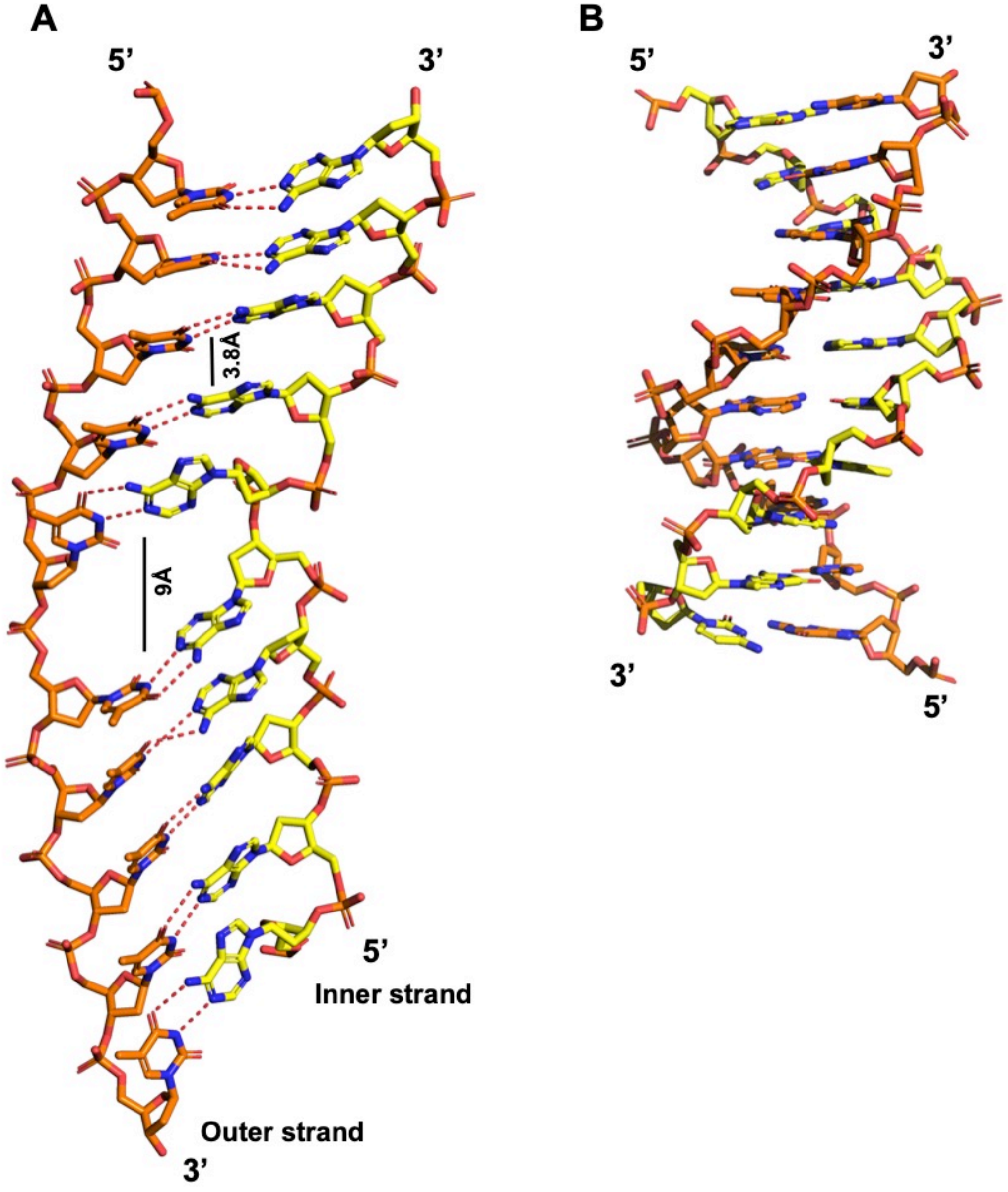


**Fig. S7. Conformation of the annealed duplex bound to LiRecT and comparison with B-form DNA.** (***A***) Segment of 10 bp from the LiRecT filament with the inner strand in yellow and the outer strand in orange. The bp stack with approximately 3.8Å spacing, except where they are separated to approximately 9Å by insertion of the β1-β2 hairpin. (***B***) 10 bp of B-form DNA, drawn to scale. Coordinates of B-DNA are from PDB code 1BNA (8). Notice that the DNA from the LiRecT filament is highly extended and under-wound, but still forms normal Watson-Crick base pairs, as indicated by the dotted lines.

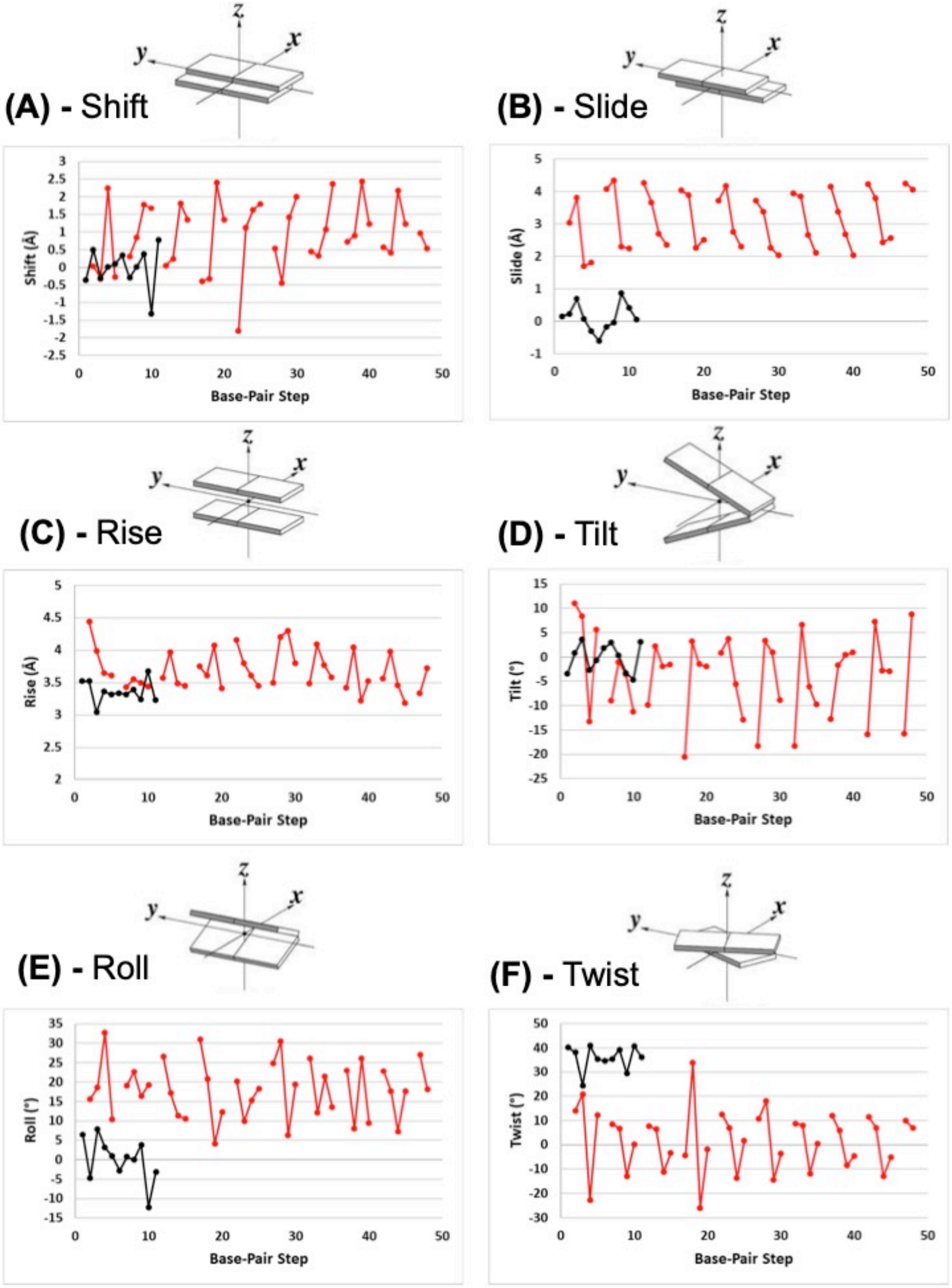


**Fig. S8. Local base-pair step parameters of the annealed duplex bound to LiRecT.** The parameters for the DNA bound to LiRecT (red lines) and B-form DNA from PDB code 1BNA (black lines) were calculated using the 3DNA program (9). Gaps in the red lines for LiRecT occur at every 5$^{th}$ base-pair step where consecutive base pairs are separated by ~9Å. The base-pair step number (n) on the x-axis indicates the base-pair step between the nth base and the (n+1)th base. Cartoons above each graph (***A-F***) show a graphical representation for each parameter (9).

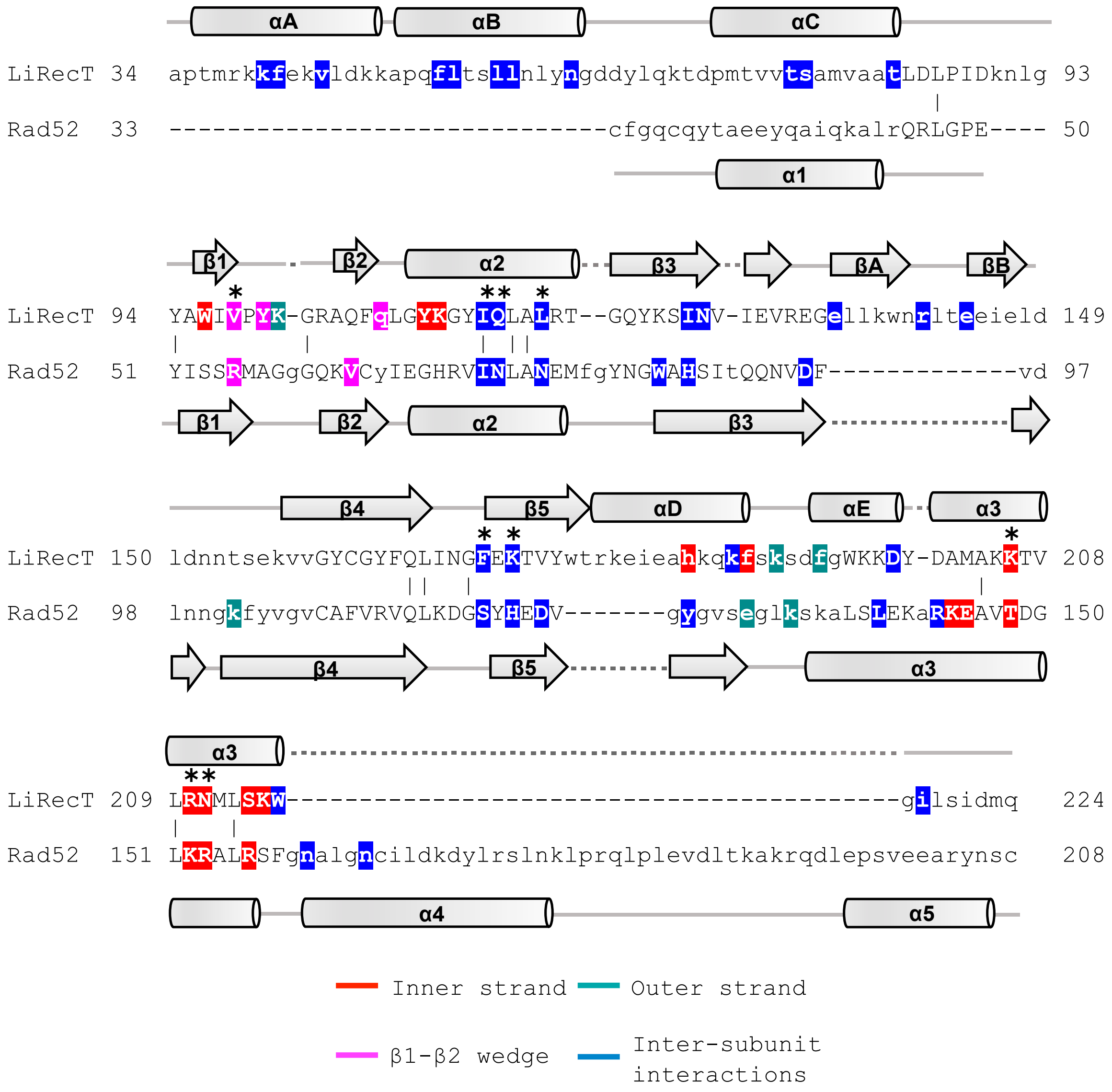


**Fig. S9. Structure based sequence alignment of LiRecT and Rad52.** Structurally equivalent residues in the alignment are in uppercase, structurally non-equivalent residues (including insertions) are in lowercase. Amino acid identities are marked by vertical bars. The shading indicates amino acids of each protein that contact the inner strand (red), outer strand (teal), form the β1-β2 hairpin wedge (violet), or form inter-subunit interactions (blue). The ∗ symbols indicate key functional residues of LiRecT that are structurally conserved with Rad52 in the alignment.

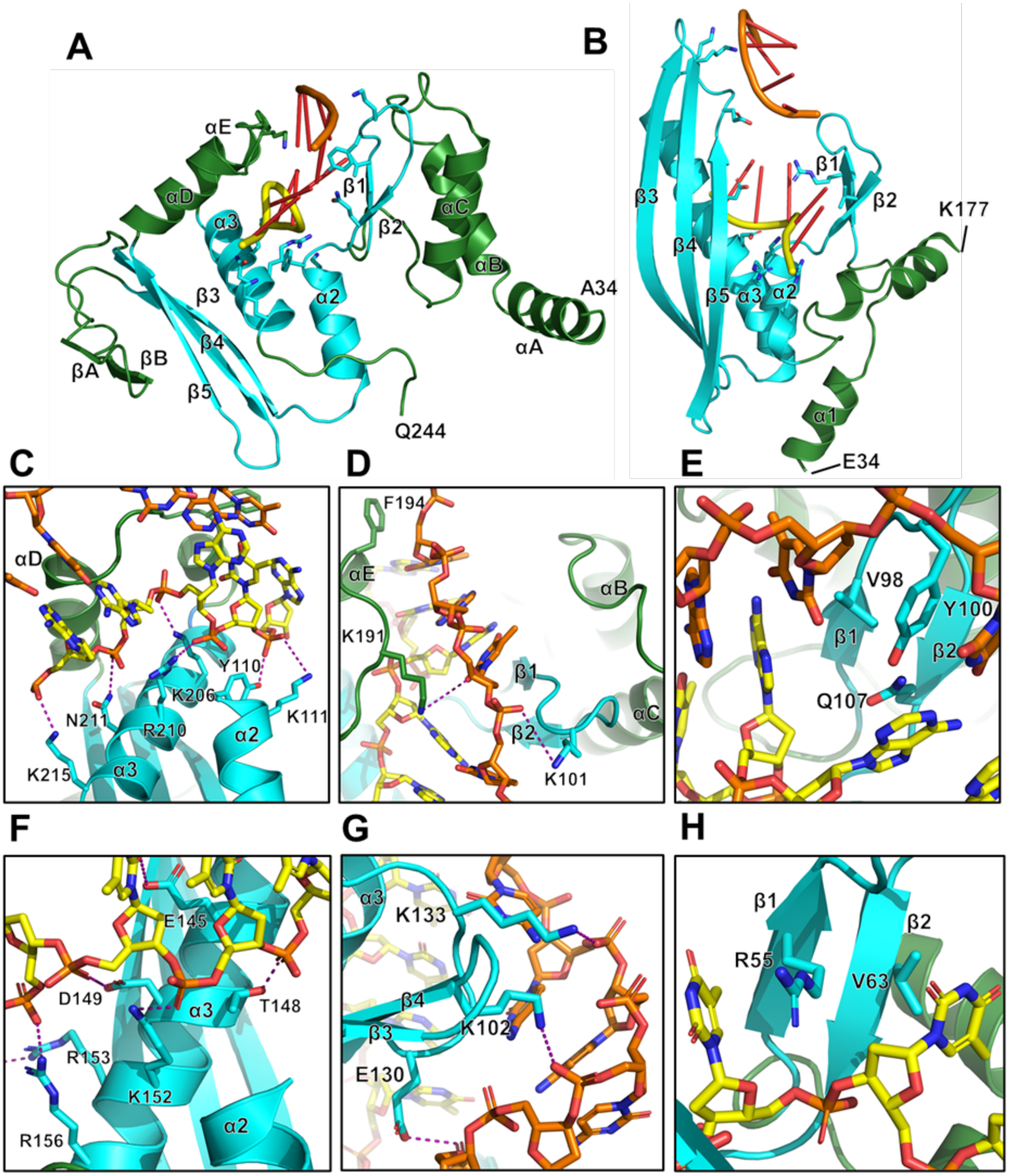


**Fig. S10. Structural homology and DNA binding residues of LiRecT and Rad52** (**A**) A monomer of LiRecT from the cryo-EM structure, with the core common to Rad52 colored in cyan. (**B**) Structure of Rad52 with DNA bound to the inner site (PDB ID 5RXZ), with the ssDNA from a separate structure with ssDNA bound to the outer site (PDB ID 5XS0) superimposed (10). (**C**) Residues in α2 and α3 of LiRecT contact the sugar phosphate backbone of the inner strand (yellow). (**D**) Residues in the two loop regions of LiRecT at the outer portion of the groove contact the outer strand (orange). (**E**) Residues from the β1-β2 hairpin that inserts into the bp at every 5$^{th}$ base pair step. (**F-H**) The three corresponding regions of interaction are shown for Rad52 (10).

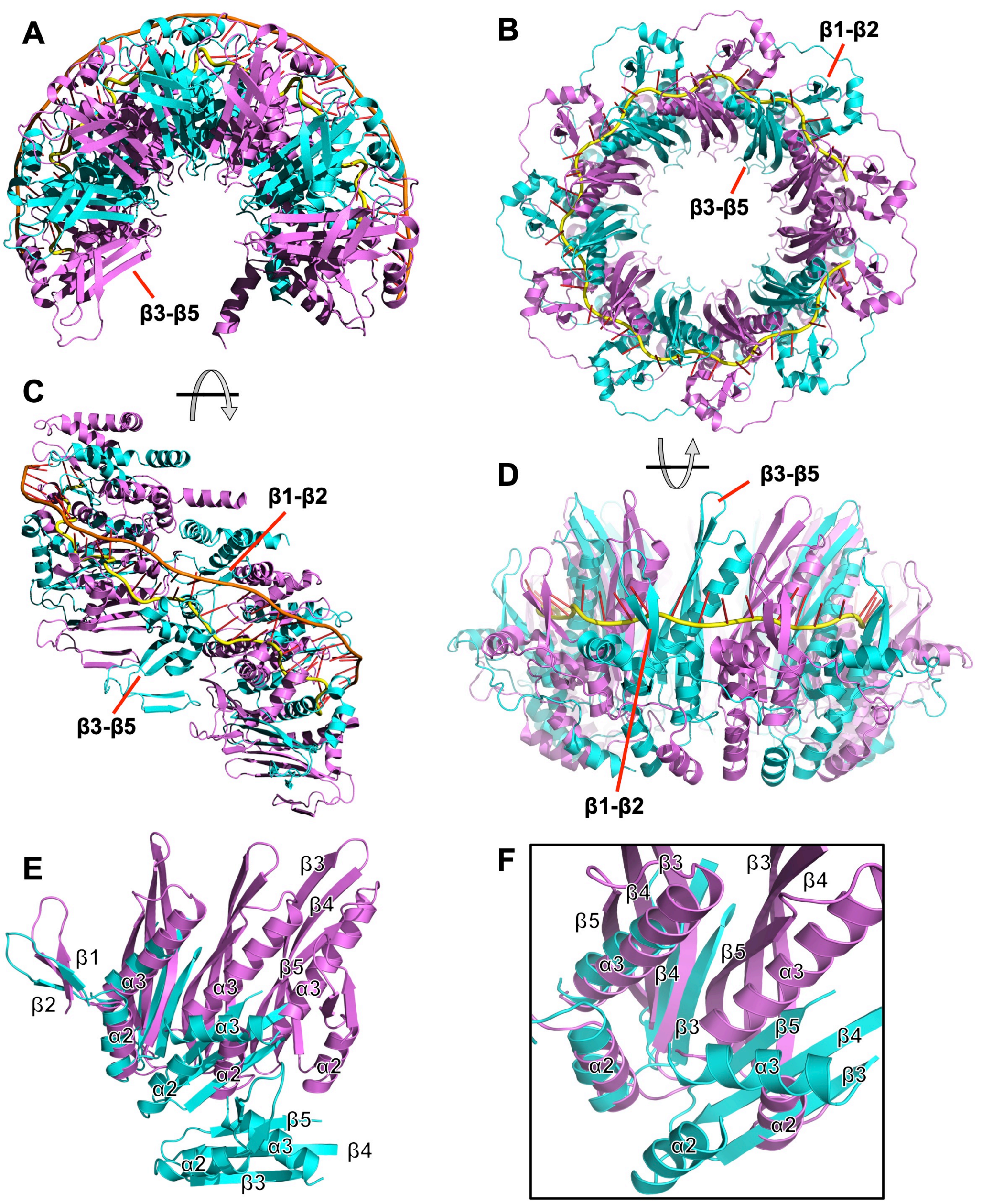


**Fig. S11. Comparison of the inter-subunit packing in LiRecT and Rad52.** (***A***) View from the bottom of the LiRecT filament looking up the helical axis. Notice that the DNA is on the outer surface of the filament and the β3–β5 sheet is on the bottom surface (facing the reader). (***B***) Top view of the closed 11-mer ring of Rad52 bound to a dT40 oligonucleotide (PDB ID 5XRZ; 10). Notice that the β3–β5 sheet is on the inner surface of the ring, and the β1–β2 hairpin is on the outer surface. (***C,D***) Side views of panels *A* and *B*, respectively. Notice in panel *C* that the β1–β2 hairpin of LiRecT is above the DNA-binding groove, and the β3–β5 sheet is below it. Notice in panel *D* that the β1–β2 hairpin is on the outside of the DNA binding groove while the β3–β5 sheet is on the inside. (***E***) The core portions of three subunits of Rad52 (violet) and LiRecT (cyan) are aligned by their left-most subunits, to show the reorientation in subunit packing to form the ring (violet) or the filament (cyan). (***F***) Close-up view of the subunit interface shows that the two proteins pack into their oligomers using the same structural elements, β3 and α2 from the left subunit and β5 and α3 from the right subunit, but in slightly different orientations.

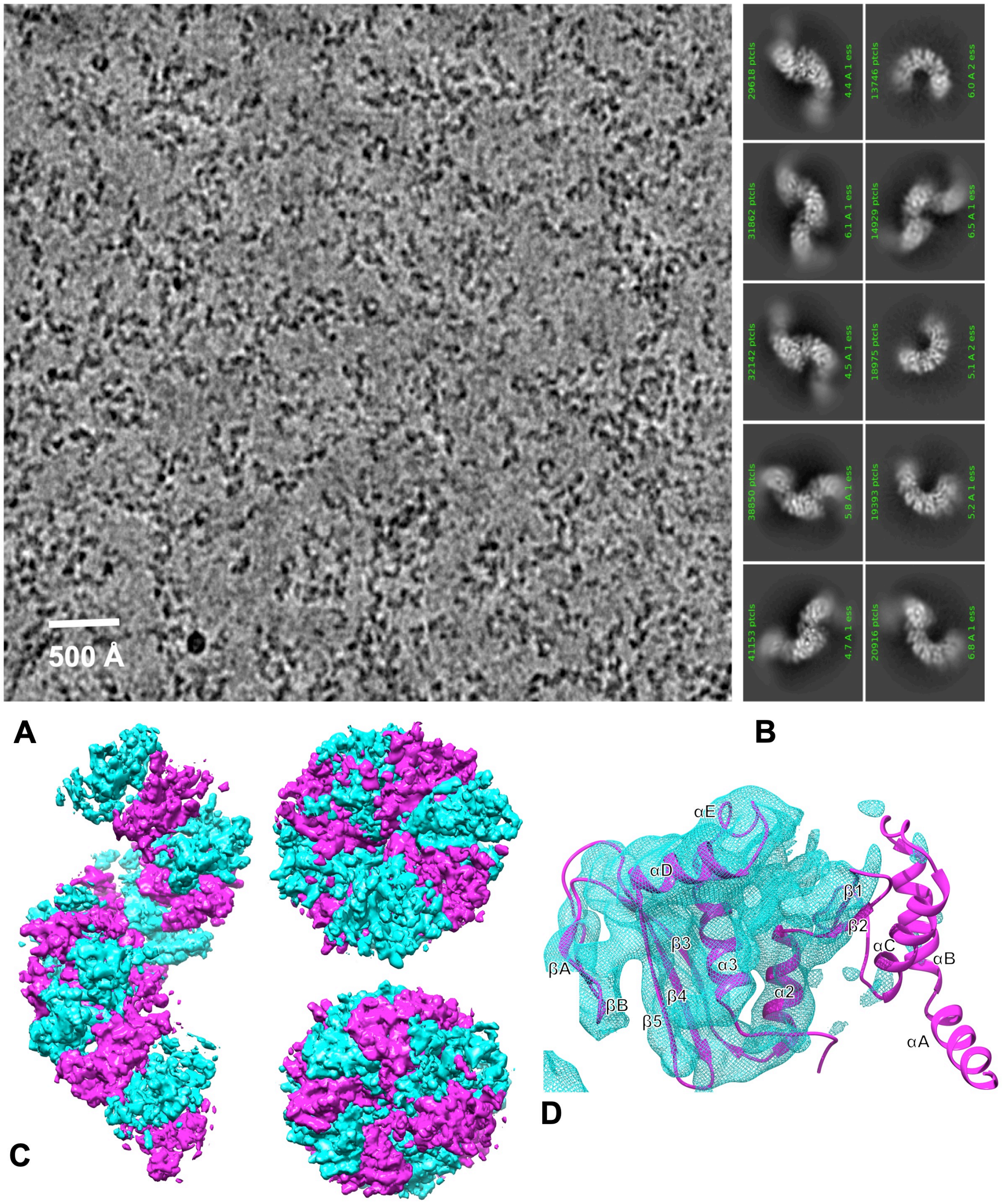


**Fig. S12. Cryo-EM analysis of LiRecT bound to 83- ssDNA.** (***A***) Krios-K3 image at 81,000x. (***B***) Selected 2D-class averages generated by cryoSPARC (3). (***C***) Views of the 5.1 Å 3D reconstruction after density modification in PHENIX. 12 subunits of LiRecT were placed into the density and rigid body refined in PHENIX. The filament with ssDNA is more tightly wound than that with annealed duplex and contains ~8 subunits per turn as opposed to ~10. (***D***) View of a monomer from the 3D reconstruction. The density for the part of the LiRecT monomer below the DNA binding groove (on the lower rim of the filament as viewed in Fig. 1 of the main text) clearly shows the positions of the β1-β3 sheet, the βA-βB hairpin, α2, α3, αD, and αE. By contrast, the absence of density for the part of the filament above the DNA binding groove (as viewed in Fig. 1) shows that the αA-αC bundle, and the β1-β2 hairpin, which form the N-terminal lobe of each LiRecT monomer, are disordered. We hypothesize that this N-terminal lobe clamps down on the DNA once a complementary strand is bound, as in the complex with annealed duplex. The DNA binding groove located above α2 and α3 shows strong density for bound DNA, but it could not be interpreted clearly enough to model.

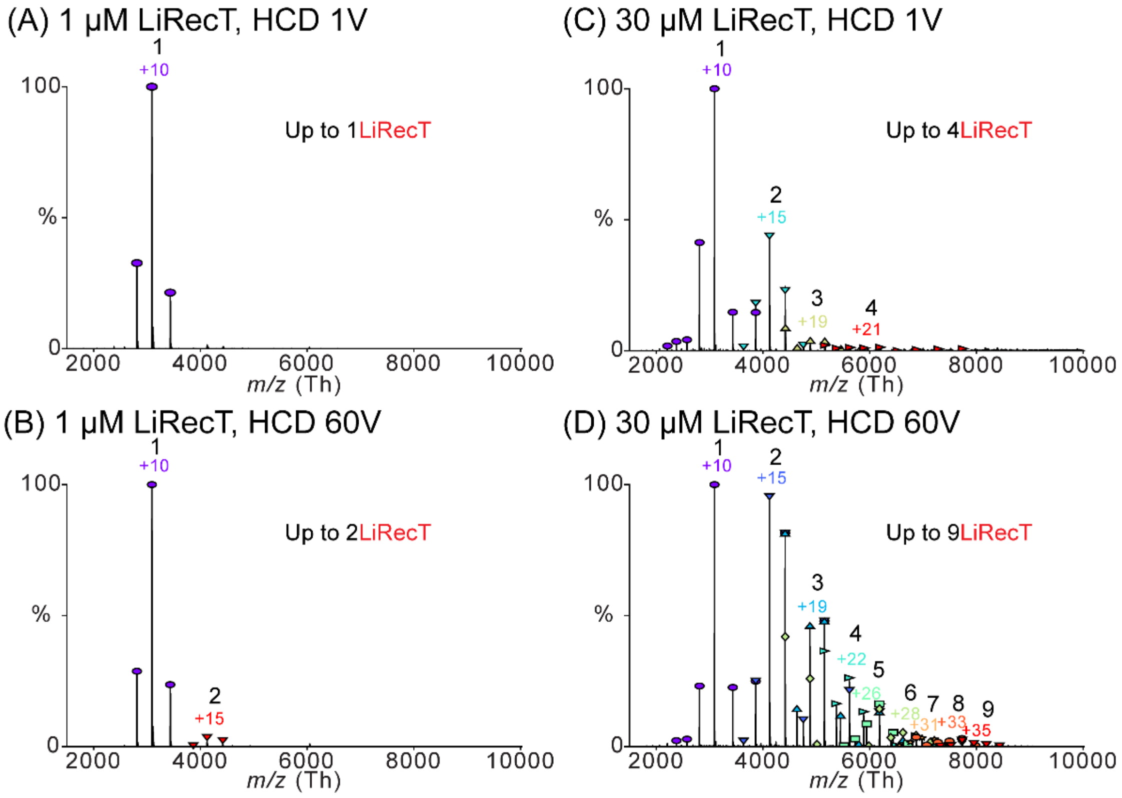


**Fig. S13. Native mass spectra of LiRecT protein alone.** Protein was dialyzed into 100 mM ammonium acetate and injected at 1 μM (A & B) or 30 μM (C & D). The effect of HCD is compared for 1 μM LiRecT with (A) 1 V HCD versus (B) 60 V HCD, and for 30 μM LiRecT with (C) 1 V HCD versus (D) 60 V HCD. Within each spectrum the charge state distribution for each species is indicated by the numbers in different colors. The number in black above each charge state indicates the LiRecT oligomeric state for each distribution.

(A) 2 µM LiRecT, HCD 60 V

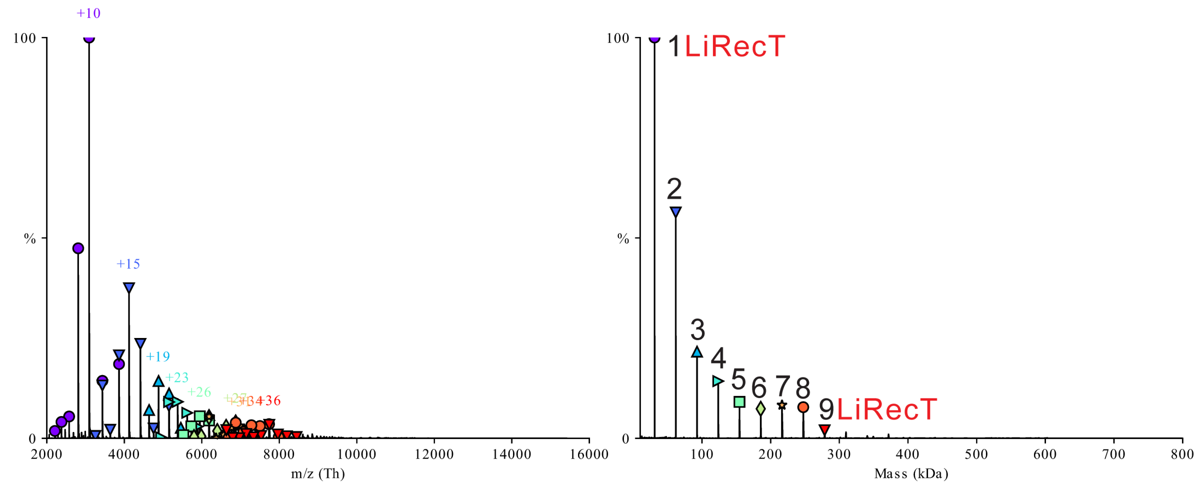


(B) 2 µM LiRecT + 5 µMnt 83-, IST 10V + HCD 90V

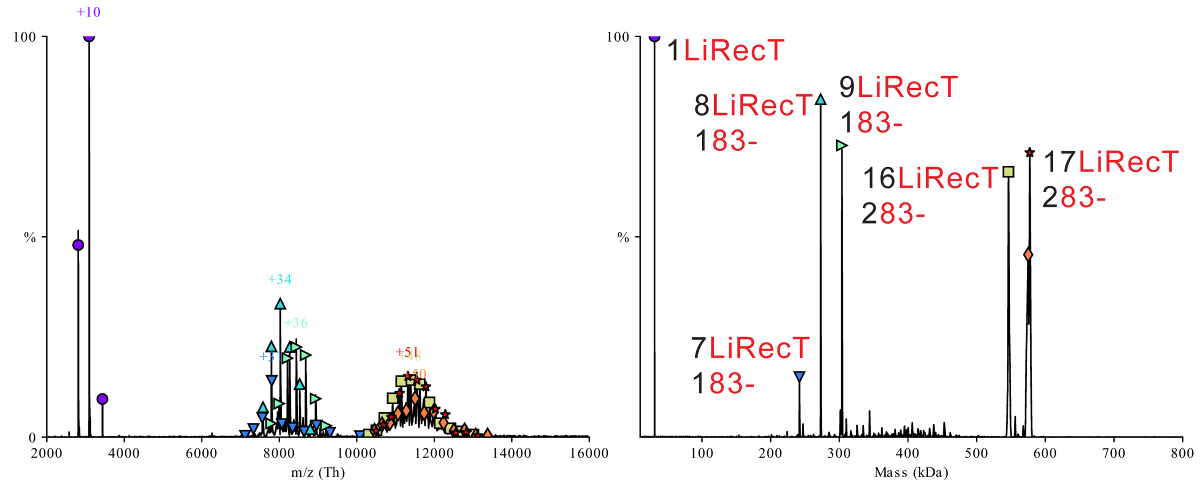


(C) 2 µM LiRecT + 5 µMnt 83+, IST 10V + HCD 90V

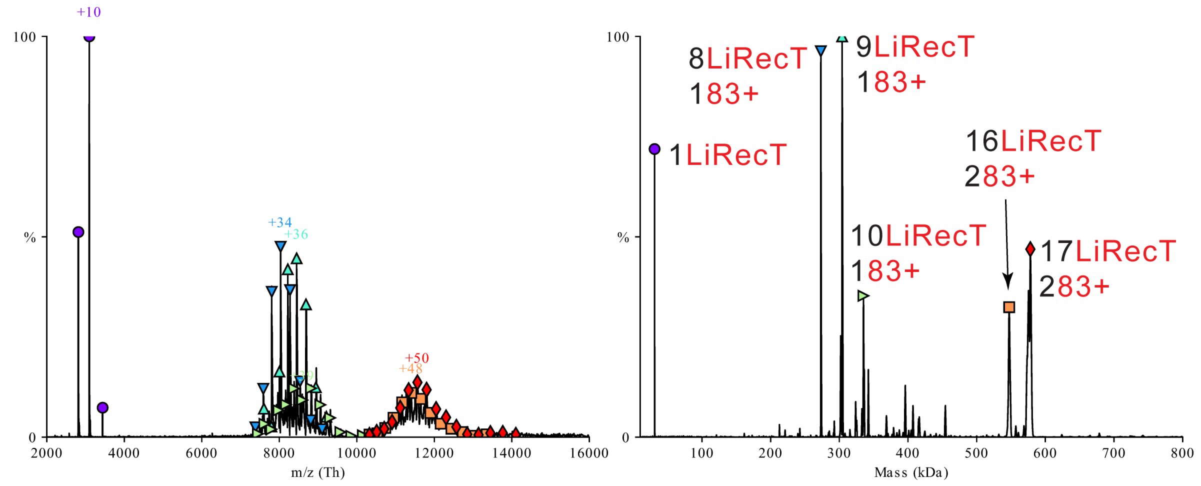

(D) 2 μM LiRecT + 5 μMnt 80-, IST 10V + HCD 90V

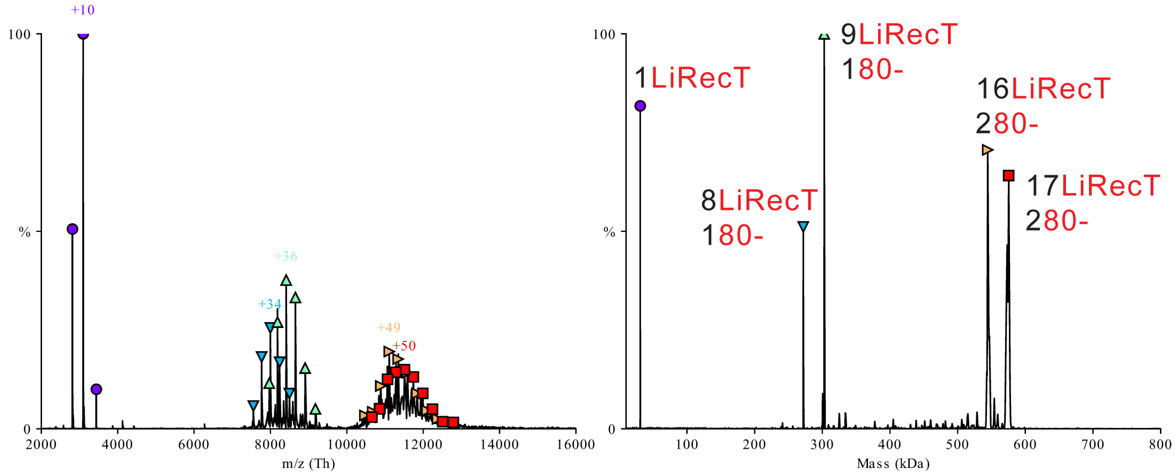


(E) 2 μM LiRecT + 5 μMnt 80+, IST 10V + HCD 90V

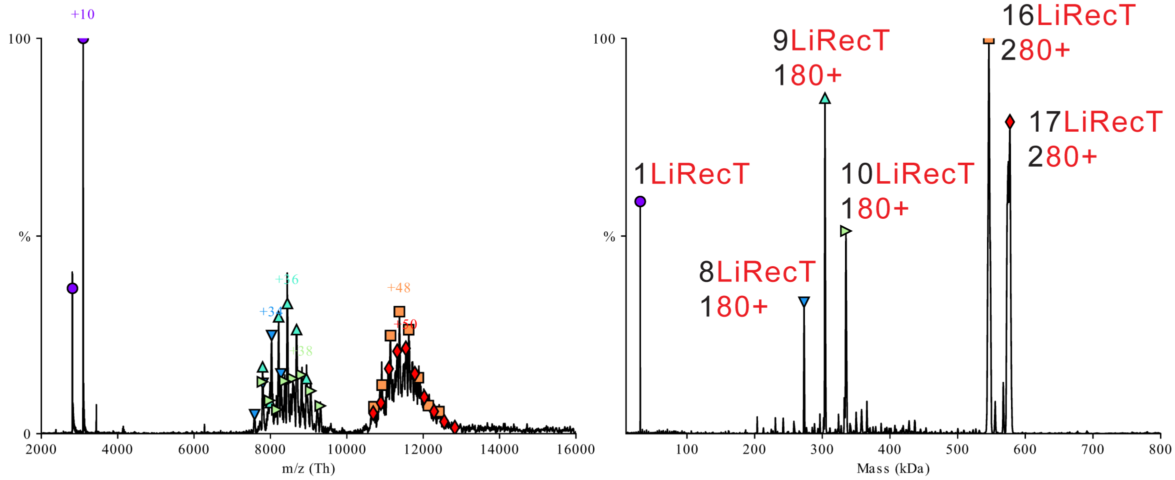


(F) 2 μM LiRecT + 5 μMnt 75-, IST 10V + HCD 90V

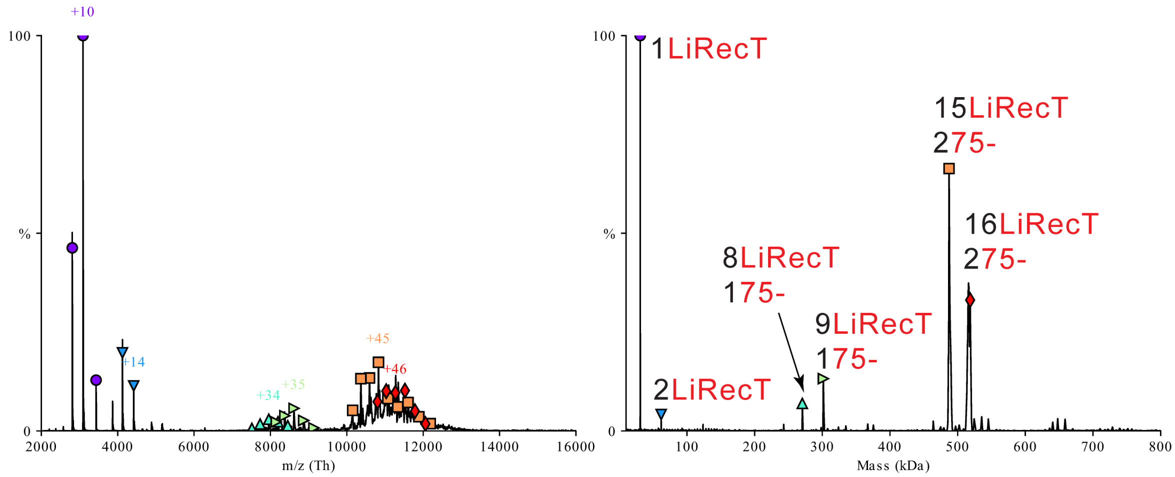

(G) 2 μM LiRecT + 5 μMnt 75+, IST 10V + HCD 90V

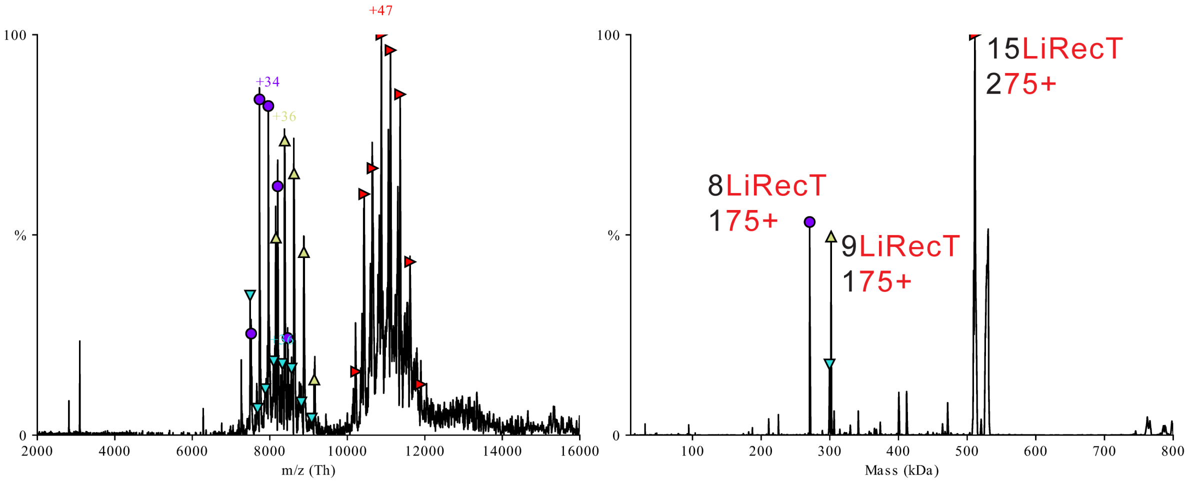


(H) 2 μM LiRecT + 5 μMnt 83-:83+, IST 10V + HCD 90V

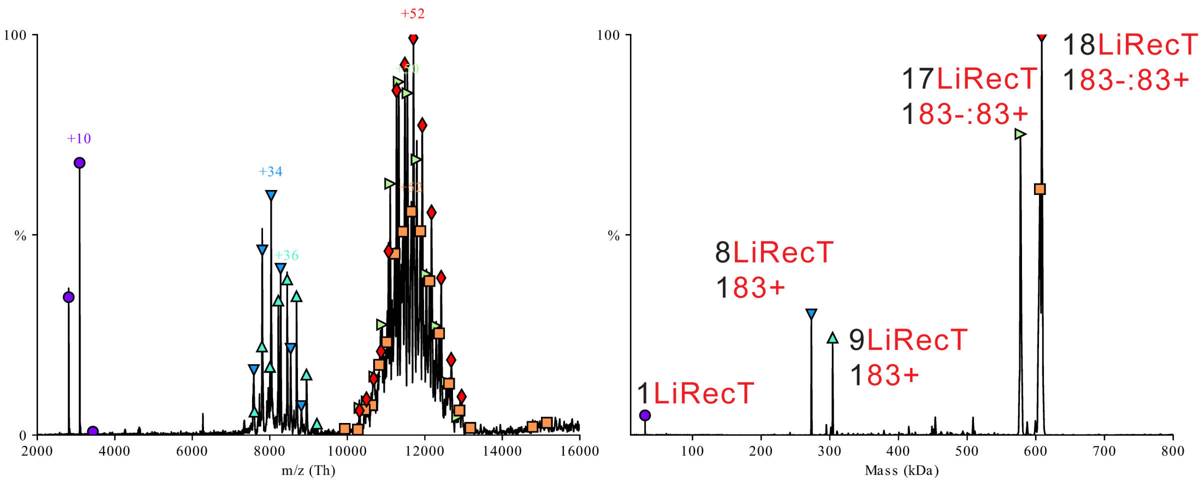


(I) 2 μM LiRecT + 5 μMnt 80-:80+, IST 10V + HCD 90V

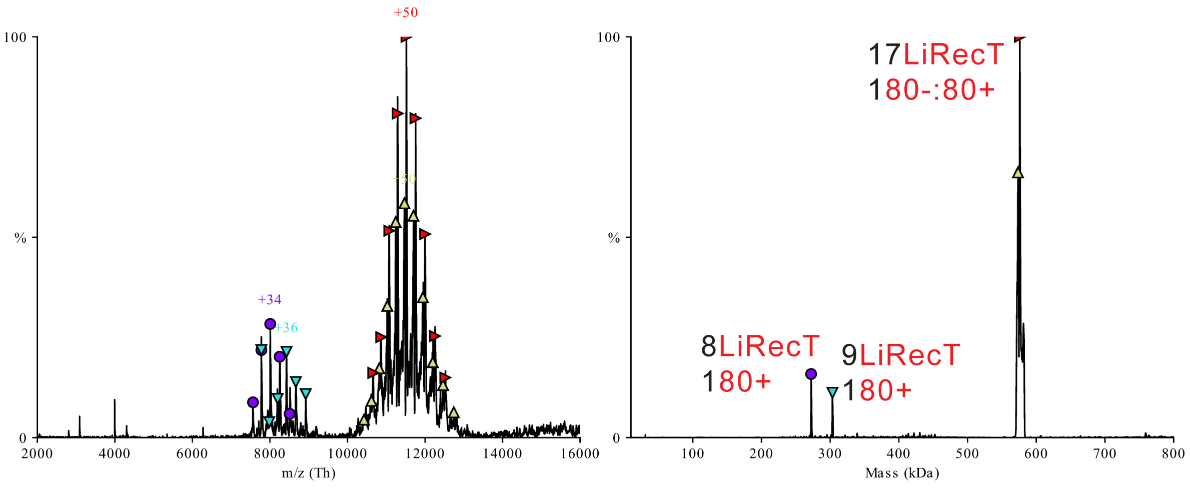

(J) 2 μM LiRecT + 5 μMnt 75-:75+, IST 10V + HCD 90V

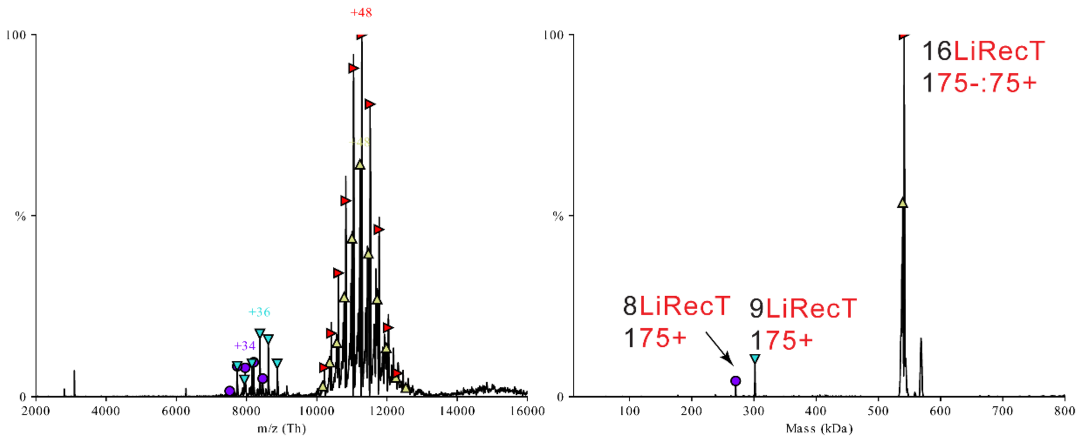


**Fig. S14.** Mass spectra (left) and zero-charge mass spectra (right) of 2 μM LiRecT mixed with different lengths and combinations of DNA . (A) LiRecT protein alone, (B) 83-, (C) 83+, (D) 80-, (E) 80+, (F) 75-, (G) 75+, (H) 83-:83+, (I) 80-:80+, (J) 75-:75+. For panels H, I, and J, two oligos were added to the protein sequentially (first:last), as described in Materials and Methods. Integrated values of the relative amounts of each species in these spectra were used to generate the data plotted in Fig. 4. All DNAs were added at a concentration of 5 μM nucleotides.

**Table S1. Cryo-EM Data Collection and Refinement Statistics[1]**

| | LiRecT + 83-:83+ (Annealed Duplex) | LiRecT + 83- (ssDNA) |
|---|---|---|
| EMDB Code | EMD-26434 | EMD-26437 |
| PDB Code | 7UB2 | 7UBB |
| Magnification | 81,000x | 81,000x |
| Voltage (Kv) | 300 | 300 |
| Dose (e-/Å$^2$) | 66 | 65 |
| Defocus range (µm) | -3.5 to -1.0 | -3.5 to -1.0 |
| Pixel size (data collection) (Å) | 0.899 | 0.4595 |
| Pixel size (reconstruction) (Å) | 0.899 | 0.899 |
| No. of movies collected | 2038 | 1619 |
| Symmetry imposed | C1 | C1 |
| Particle Images | 391,275 | 180,965 |
| Resolution (0.143 FSC, tight mask) (Å) | 3.4 | 4.5 |
| r.m.s. deviation bond lengths (Å) | 0.002 | --- |
| r.m.s. deviation bond angles (Å) | 0.515 | --- |
| MolProbability Score | 1.23 | --- |
| Clashscore | 4.49 | --- |
| C-beta outliers (%) | 0.00 | --- |
| Rotamer outliers (%) | 0.60 | --- |
| CaBLAM outliers (%) | 0.53 | --- |
| Ramachandran favored (%) | 98.94 | --- |
| Ramachandran allowed (%) | 1.06 | --- |
| Ramachandran outliers (%) | 0.00 | --- |

[1]The structure with ssDNA was limited to rigid body refinement

**Movie S1 (separate file)**.
Movie “LiRecT MoviePresentation” available at:
https://osf.io/nzur8/?view_only=5c76334408b543369a27406083b022f0

The movie was generated in PyMOL (5). Captions to the movie were inserted using Adobe Premier, with the following transcript:

“This movie shows seven subunits of LiRecT protein bound to 31 bp of DNA (5 bp/monomer).

The DNA is bound in a highly extended and under-wound conformation that appears to be a novel duplex intermediate of DNA annealing.

The complex was prepared by mixing LiRecT with two complementary strands of ssDNA that were added to the protein sequentially.

The yellow strand, which we call “inner” is almost certainly the first strand added. It is bound deep in a groove with its bases exposed for homology recognition.

Several side chains of the protein contact the sugar phosphate backbone of the inner strand, and hold it in an irregular conformation that is periodically kinked.

The orange strand, which we call “outer”, forms fewer contacts with the protein, and is held in place primarily by Watson-Crick base pairs with the inner strand.

While most of the contacts are to the sugar-phosphate backbone of the DNA, a beta-hairpin of the protein wedges into the base pairs to separate them, at every 5 bp step.

Neighboring subunits of LiRecT bind to one another via two different sets of interactions.

The first set, above the DNA binding groove, involves the N-terminal helix bundles, which form a core of hydrophobic interactions surrounded by a few ion pairs.

A second set of interactions, below the DNA binding groove, forms a similar hydrophobic core using the three-stranded beta-sheet and two central alpha helices.

We hypothesize that the N-terminal helix bundles clamp down on the duplex after the second strand is added, to stabilize the complex and consolidate annealing.”
